# Cross regulation between the molecular clock and kidney inflammatory, metabolic and fibrotic responses

**DOI:** 10.1101/2022.05.18.492458

**Authors:** Carlos Rey-Serra, Jessica Tituaña, Terry Lin, J. Ignacio Herrero, Verónica Miguel, Coral Barbas, Anna Meseguer, Ricardo Ramos, Dmitri Firsov, Amandine Chaix, Satchidananda Panda, Santiago Lamas

**Author notes:** **Correspondence:** Dr. Santiago Lamas, Program of Physiological and Pathological Processes, Centro de Biología Molecular Severo Ochoa (CSIC-UAM), Nicolás Cabrera 1, Campus UAM, 28049 Madrid, Spain.

## Abstract

Chronic kidney disease is a highly prevalent condition that remains a major clinical and biomedical challenge. Tubulo-interstitial fibrosis is the common pathological substrate for many causes that lead to chronic kidney disease. It is characterized by profound derangements in metabolic and inflammatory responses, whereby functional tissue is replaced with extracellular matrix, leading to the suppression of renal function. Perturbations in the circadian rhythm have been associated with many human pathologies, including renal disease. However, the role of the molecular clock in the instauration of fibrosis remains incompletely understood. We investigated the relationship between the molecular clock and renal damage in experimental models of injury and fibrosis (UUO, FAN and adenine toxicity), employing genetically-modified mice with selective deficiencies of the clock components Bmal1, Clock and Cry. We found that UUO induced a marked increase in the expression of Bmal1. In human tubular epithelial cells, the pro-fibrotic mediator, TGF-β, significantly altered the expression of core clock components. We further observed that the absence of Cry drastically aggravated kidney fibrosis, while both Cry and Clock played a role in the neutrophil and macrophage mediated inflammatory response, respectively. Suppression of Cry1/2 was associated with a major shift in the expression of metabolism-related genes, underscoring the importance of metabolic dysfunction in fibrosis. These results support a reciprocal interaction between the circadian clock and the response to kidney injury.

**Translational statement:** Chronic kidney disease (CKD) is a highly prevalent clinical syndrome that still poses major clinical challenges. Kidney fibrosis underlies many cases of CKD and therapies against it are of very limited efficacy. Alterations in circadian rhythms (CR) are relevant in patients with CKD, but very little is known about the relationship between CKD and CR. Our study shows that disruption of the molecular clock can impact kidney inflammation and fibrosis and that, reciprocally, kidney fibrosis can alter the expression of clock components. A better understanding of this crosstalk could open new therapeutic avenues for the prevention and treatment of CR-related CKD.

## Introduction

Organ fibrosis is a primordial consequence of chronic inflammatory disease characterized by an excessive accumulation of extracellular matrix proteins (ECM). In the kidney, fibrosis is the ultimate stage of cellular responses to chronic inflammation and the pathological convergent substrate for several entities of different origin, leading to chronic kidney disease (CKD)^1^. Whereas the understanding of the molecular and cellular basis of fibrosis, including that of the kidney, has experienced substantial progress in the last decade^2–4^, options for prevention or treatment are scarce, with only a few drugs offering limited therapeutic advantage in the case of idiopathic pulmonary fibrosis^5^. Renal tubulo-interstitial fibrosis is characterized by a profound alteration of metabolism within the tubular epithelial cell, majorly related to a drastic reduction in fatty acid oxidation (FAO)^6^. We recently showed that genetically mediated FAO enhancement in the renal tubule resulted in significant protection from experimental fibrosis of different origins^7^. We also found that levels of the FAO-related mitochondrial enzyme CPT1a were reduced in patients with CKD and that levels of long and middle chain acylcarnitines were significantly altered. Thus, kidney fibrosis should now be envisioned as an example of metabolic failure, whereby inflammation and mitochondrial dysfunction contribute to perpetuate a vicious cycle that results in extracellular matrix deposition, fibrosis and establishment of CKD.

Many physiological functions of most tissues and cells across living organisms exhibit daily periodic fluctuations related to the circadian rhythm (CR), majorly conditioned by predictable environmental cues such as the light/dark cycle. The CR is composed by a master clock, located in the hypothalamus, which is entrained by light and can synchronize peripheral clocks via neuronal and humoral signals^8^. At the molecular level, the peripheral clocks are regulated by a transcriptional feedback loop in which clock components such as Arntl1 (Bmal1) and Clock activate the transcription of their own repressors including Per and Cry^9^. Perturbations in the CR, including the molecular clock, have been associated with many pathologies including renal disease^10–12^. Additionally, there is an important crosstalk between the circadian clock and metabolism^13^. However, the role of the molecular clock in the development of kidney fibrosis remains largely unknown. In this study, we explored the relationship between the molecular circadian clock and the inflammatory and fibrotic responses in several experimental renal injury models. UUO-induced kidney damage was associated with a significant change in the expression of Bmal1, consistent with the action of TGF-β in human tubular epithelial cells. We found that selective deficiencies in the core clock components Bmal1, Clock and Cry were associated with differential responses regarding inflammation and fibrosis. Strikingly, the absence of Cry 1/2 promotes a major shift in the expression of metabolism-related genes and markedly aggravates the fibrotic phenotype.

## Methods

### Genetically modified mouse models in the circadian clock (clock mutant mice)

Clock^Δ19^ mice, Per2::Luc mice (kindly provided by Dr. Takahashi’s lab, Texas, USA), Bmal1^lox/lox^/Pax8-rtTA/LC1 (Bmal1 cKO), Bmal1^lox/lox^/CRE-ER (Bmal1 KO) and Cry1/Cry2 mutant mice (double KO mice (CDKO), Cry1 KO, Cry2 KO and double heterozygous (CDHet)) were bred on the genetic background of the C57BL/6J mouse and were generated and characterized as previously described^14–17^. Clock^Δ19^ mice were crossed with Per2::Luc mice, using the Per2::Luc strain as control (WT), Bmal1^lox/lox^ mice (WT) were used as control for the Bmal1 cKO and the Bmal1 KO and male C57BL/6J mice (WT) purchased from The Jackson Laboratory (Bar Harbor, ME, USA) were used as controls for the CDKO mice. The genotype of the animals was confirmed by PCR by using the primers listed in **Table S1**. The animals were maintained *ad libitum* on regular chow diet, under constant temperature and humidity in a 12-hour light/dark cycle. Doxycycline and Tamoxifen were used for the induction of Cre expression in 8 weeks-old Bmal1 cKO and Bmal1 KO mice, respectively. Mice were treated daily with 2 mg/ml of doxycycline in drinking water ^17^ or 3.7 mg of tamoxifen administered by oral gavage for 2 weeks or 5 days, respectively. Subsequent experiments were performed four or two weeks after the end of the treatment, respectively, in order to avoid potential side effects related to their administration. Mouse activity was tested in the Bmal1 KO and their WT mice after Cre expression induction with tamoxifen using the Mouse Home Cage Running Wheel (The Columbus Instruments, OH, USA).

### Mouse models of chronic kidney disease

<u>Unilateral Ureteral Obstruction (UUO)</u> procedure was performed as described previously^18^. Briefly, 12-14 weeks old mice were anesthetized with 2% isoflurane. The hair in the abdominal area was shaved. An incision was made in the abdominal wall to expose the left kidney, the ureter was ligated twice and severed between the two ligatures and the kidney was returned gently to its place. Finally, the abdominal incision was closed with sutures and buprenorphine was used as an analgesic. <u>Folic acid nephropathy (FAN)</u> was carried out as previously described^19^. Mice were given 250 mg/kg of folic acid intraperitoneally (i.p) (Sigma-Aldrich) resuspended in 0.3 M sodium bicarbonate (vehicle) and control mice were given 0.1 ml of the vehicle. For <u>adenine- induced renal failure (ADN)</u>, mice were given 50 mg/kg of adenine in 0.5% carboxymethyl cellulose (CMC) (Wako Pure Chemical Industries Ltd., Osaka, Japan) daily by oral gavage as described previously^20^. Control mice were given 0.1 ml of 0.5% CMC (vehicle). Mice were sacrificed 3, 7, 15 or 25 days after UUO; 7 and 15 days after FAN or 25 days after ADN. The three procedures and the sacrifice of the mice were performed at ZT9 (where ZT0 means light on and ZT12 means light off). Blood samples were collected right after the sacrifice by cardiac puncture and kidneys were collected after perfusion with PBS. Number of mice described in **Table S2**. **Cell culture.** Immortalized renal human proximal tubule epithelial cells RPTEC/TERT1 (HPTEC) were obtained from the American Type Culture Collection (ATCC®; #CRL-4031). These were cultured in DMEM/F12, GlutaMAX supplement (Dulbecco’s modified Eagle’s medium 1:1 (v/v)) (Life technologies, MA, USA) supplemented with 20 mM Hepes, 2% (v/v) fetal bovine serum (FBS) (HyClone Laboratories, Logan, UT), 5 *μ*g/ml Apo-transferrin (Sigma-Aldrich, St. Louis, MO, USA), 5 *μ*g/ml Human Insulin Solution (Sigma), 50 nM dexamethasone (Sigma), 3 nM 3,3’,5-Triiodo-L-thyronine sodium salt (Sigma), 10 ng/ml EGF (Sigma), 60 nM Selenium (Sigma), 50 units/ml penicillin and 50 *μ*g/ml streptomycin (Gibco, Rockville, MD, USA) at 37°C and 5% CO_2_. Sub-culturing was performed every 3-5 days. Other cell lines and i*n vitro* experiments were respectively cultured and performed as described in supplementary methods.

### Statistical analysis

Data analysis were performed using GraphPad Prism 8.0 (GraphPad Software, La Jolla, CA, USA). Data are represented as mean ± standard error of mean (SEM). Statistical differences were determined with the non-parametric Mann-Whitney test when two independent groups were analyzed, while two-way ANOVA and Tukey’s post-test were used for more than two independent groups and two independent variables and the nonparametric Wilcoxon test was used for paired analysis. A P-value of 0.05 was considered to be statistically significant (^*/#^P<0.05, ^**/##^P<0.01, ^***/###^P<0.001).

## Results

### The expression of Bmal1 and other clock components is differentially altered in three different models of kidney fibrosis

To investigate the influence of kidney fibrosis on the expression of core clock components of the circadian machinery we determined the gene expression levels of Bmal1, Bmal2, Clock, Npas2, Cry1, Per1, Per2, Cry2, Nr1d1 and Nr1f1 in the UUO, ADN and FAN models^7^ (Fig. 1A). We found that after UUO, the expression of Bmal1 was increased at all time points analyzed in comparison with their contralateral kidneys. Interestingly, its expression progressively increased, peaking 25 days after UUO, in good correlation with the mRNA levels of the profibrotic cytokine TGFβ and the fibrosis-related genes Fn1 and Col1a1 (Fig. 1B). This pattern was also consistent in the other two models, at 7 and 15 days after FAN and at 25 days after ADN (Fig. 1C, D). The analysis of the mRNA levels of other clock components revealed that most of the clock genes were also differentially expressed after UUO and FAN compared to control kidneys (Fig. S1A, B), while only Nr1d1 expression was significantly altered 25 days after ADN (Fig. S1C). Of note, the UUO model presents higher expression levels of Bmal1 than the FAN and ADN models for the same time point, a pattern that is maintained with levels of TGFβ, Col1a1, Fn1 and collagen deposition (Fig 1B-D; Fig. S1D, E). This observation, together with the greater variability observed in FAN and ADN due to a higher mortality rate associated to these models, pointed to the UUO as the most reliable model for further analysis. These results indicate that kidney injury elicits significant changes in the expression of several components of the molecular clock.

**Figure 1:**
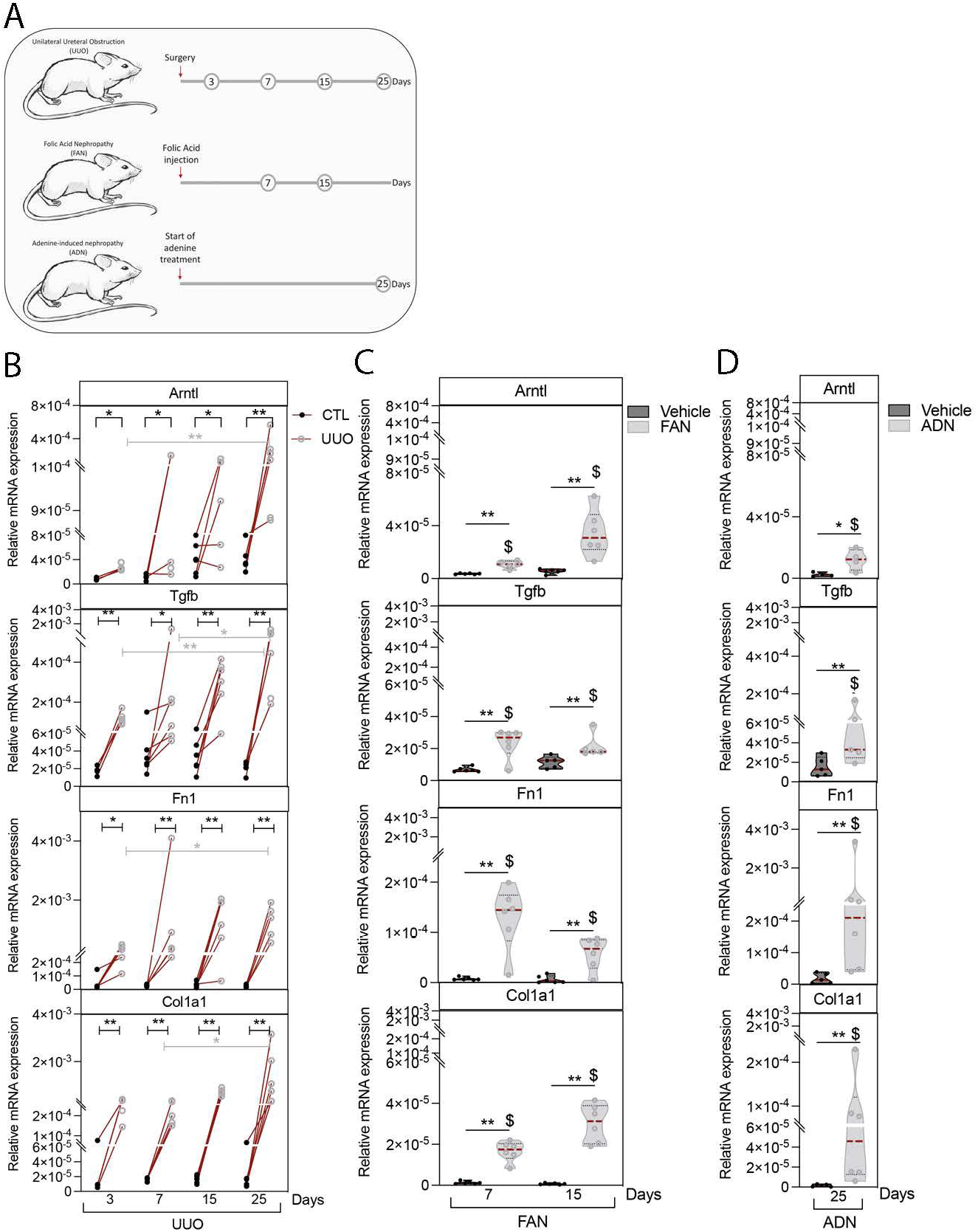
The gene expression or Arntl and fibrosis-related genes is upregulated in kidney fibrosis. (A) Schematic of experimental design of WT mice subjected to UUO for 3, 7, 15 and 25 days, FAN for 7 and 15 days and ADN for 25 days. (B) mRNA expression of Arntl, Tgfb, Col1a1 and Fn1 in kidneys from mice subjected to UUO for 3, 7, 15, 25 days (n=6 per condition), data are represented as individual values. Each obstructed kidney was linked to their corresponding contralateral. (C) mRNA expression of Arntl, Tgfb, Col1a1 and Fn1 in kidneys from mice subjected to FAN for 7 and 15 days (n= 6 mice per condition), (D) mRNA expression of Arntl, Tgfb, Col1a1 and Fn1 in kidneys from mice subjected to ADN for 25 days (n=5 mice per condition). CTL: Contralateral. Data are represented with violin plots as the median with interquartile range. ^*^P<0.05, ^**^P<0.01 compared to control kidney (black asterisk) or compared to a different time point for each gene within the same model (grey asterisk). ^$^P<0.05 compared to the UUO model at the same time point.

### TGFβ upregulates the expression of molecular clock components in human proximal tubular epithelial cells (HPTEC)

TGFβ, the archetypal profibrotic cytokine is significantly involved in kidney fibrosis in humans^21^. Thus, we evaluated its effect on clock components expression in an immortalized cell line of human proximal tubular epithelial cells (HPTEC). As expected, TGFβ increased the expression of the fibrotic markers FN1 and Col 1a1 in HPTEC (Fig. 2 A). Interestingly, treatment of HPTEC with TGFβ augmented the expression of Bmal1, Bmal2, Clock and Cry1 in a time-dependent fashion, while the expression of the rest of the clock components analyzed was not modified (Fig. 2 B-D). TGFβ-mediated increased in Bmal1 and Cry1 expression was abolished by treatment with SB505124, a TGFβ1 receptor antagonist, as demonstrated by qPCR, western blot and immunofluorescence microscopy (Fig. 2 E-I), thus attesting to the mediation of TGFβ signaling by Alk 4-5-7 receptors. These data correlate with the results obtained in primary kidney cells from C57BL6/J wild type mice (Fig S2A) and in human primary PTEC (HprimPTEC) (Fig S2B-D). Moreover, our data revealed that Bmal1 is the clock component showing the earliest upregulation, increasing just 4 hours after TGFβ treatment (Fig. 2 B). Furthermore, the analysis of Bmal1 pre-mRNA levels revealed an increase in their expression in fibrotic kidneys 25 days after UUO and in HprimPTEC after TGFβ1 treatment (Fig S3A-C), suggesting a potential transcriptional mechanism. *In silico* evaluation of the Bmal1 gene, Arntl, using the R package for JASPAR2018^22^, revealed two potential binding sites for Smad3/4 with a relative score of 88/96% and 83%, respectively (Fig. S4A-C). These sites belong to a distal enhancer located at 8376 bp upstream from the transcription start site (Fig. S4A) and are highly conserved among vertebrates (Fig. S4D). However, overexpression of Smad3 did not result in an increase of Bmal1 expression (Fig. S4E, F). In keeping, the analysis of transcriptional activity in the presence of plasmids bearing the enhancer regions corresponding to the Smad3 binding sites described above did not result in a significant increase (Fig. S4G). While this is compatible with a Smad3-independent regulation of Bmal1, the fact that the expression of Col1a1 (Fig.S4E) and the phosphorylation of Smad3 were not affected by Smad3 overexpression (Fig. S4F) implies that the latter is not sufficient to activate RSmad-dependent TGFβ1 signaling and hence Smad3-dependent regulation of Bmal1 cannot be completely excluded.

**Figure 2:**
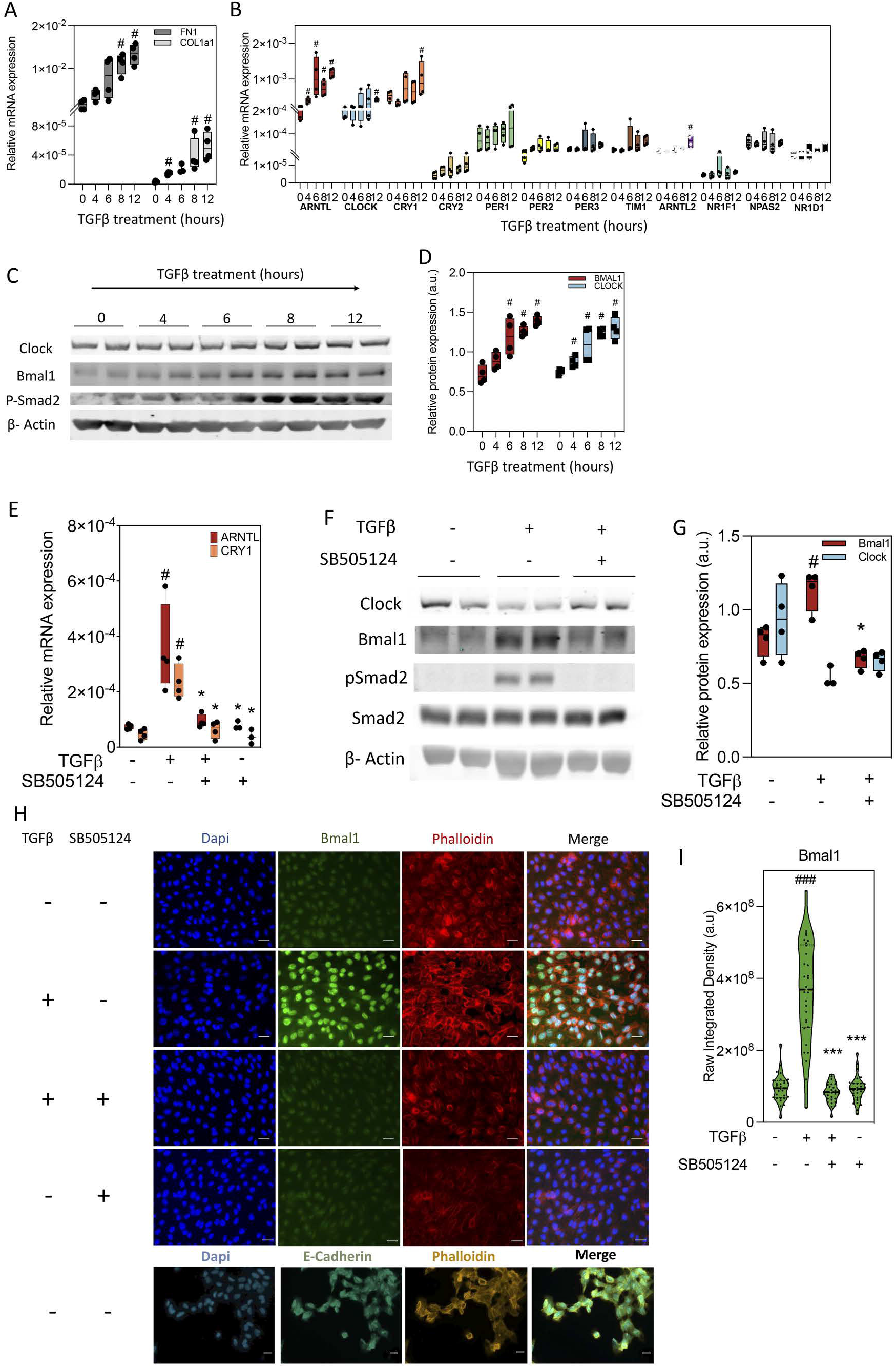
The expression of circadian oscillators is upregulated by TGF-β in proximal tubular epithelial cells. (A-D) Evaluation of the expression of circadian oscillators in synchronized HPTEC cells treated with TGF-β for 0, 4, 6, 8 and 12 hours. (A) Relative mRNA levels of fibrosis-related genes, data are represented with box plots as the median with interquartile range (n=4 per condition); (B) Relative mRNA levels of circadian oscillators, data are represented with box plots as the median with interquartile range (n=4 per condition); (C) immunoblots showing the Bmal1, Clock and pSmad2 expression. β-actin was used for normalization; (D) Relative protein expression of Bmal1 and Clock obtained by densitometry of images from C and normalized with β-actin, data are represented with box plots as the median with interquartile range (n=4 per condition); (E-I) Evaluation of the expression of Bmal1, Cry1 and Clock in synchronized HPTEC cells treated with TGF-β for 24 hours in the presence or absence of the selective inhibitor SB505124. (E) Relative mRNA levels of Bmal1 and Cry1, data are represented with box plots as the median with interquartile range (n=4 per condition); (F) Immunoblot depicting the expression of Clock, Bmal1, pSmad2 and Smad2. β-actin was used for normalization; (G) Relative protein expression of Bmal1 and Clock obtained by densitometry of images from F and normalized with β-actin, data are represented with box plots as the median with interquartile range (n=4 per condition); (H) Immunofluorescence microscopy of Bmal1, Dapi was used for the staining of the nucleus and Phalloidin for the actin cytoskeleton. E-Cadherin was used as epithelial marker control. Scale bar: 50 μm; (I) Quantification of the fluorescence of Bmal1 calculated from ten different fields per sample. The experiment was repeated three times and the data are represented with box plots as the median of the integrated density of all fields analyzed (a total of 30 fields per condition) with interquartile range. ^#^P<0.05, ^###^P<0.001 compared to untreated cells, ^*^P<0.05, ^***^P<0.001 compared to TGF-β-treated cells.

### Genetic deficiency of the clock component Cry aggravates UUO-induced fibrosis, while Bmal1 deficiency and Clock mutation do not exhibit significant differential impairment

To assess the contribution of individual clock components to kidney fibrosis, we performed the UUO model for 3 or 7 days in wild type (WT) mice and in two conditional mouse models with a genetic deficiency for Bmal1, the first one with a global loss of the gene (Bmal1 KO) and the second one with a loss of Bmal1 in renal tubular epithelial cells (Bmal1 cKO), Clock^Δ19^, Cry1, Cry2 and in the Cry1 and Cry2 double KO (CDKO) mice (Fig. S6A, Fig. S7A, Fig. 3A). As expected, global Bmal1 KO mice showed alterations in their activity patterns under constant darkness (Fig. S5A) and the expression of the different clock components was altered in kidneys from all mutant mice compared to their respective wild type counterparts (Fig. S5 B-E). We found that 7 days after UUO, the global absence of Bmal1 resulted in significantly less collagen deposition as documented by histological analysis (Sirius Red staining) and SMA protein expression (Fig S6B, C, E, F). However, no significant differences were observed in terms of Kim1 protein expression (Fig S6B, D), kidney function as measured by BUN and blood creatinine levels (Fig. S6G, H) and mRNA levels of the fibrotic markers Acta2, Col1a1 and Fn1 (Fig S6I). The same experimental approach was used in mice with tubule-specific deletion of Bmal1 (Bmal1 cKO) and with Clock^Δ19^ mutation (Fig S6A, S7A). Clock^Δ19^ mice showed a significant decrease in Kim1 expression (Fig. S6B, D), but no other remarkable differences were seen in either Clock^Δ19^ (Fig S6B, C, J-L) or Bmal1 cKO mice (Fig S7B-F), in terms of fibrotic phenotype, compared to their WT counterparts. By contrast, we found that the combined absence of the two Cry genes (CDKO) was associated with a more severe histological phenotype, observed by a significant increase of Kim1 expression (Fig. 3B, C). Sirius red staining revealed an upward trend of collagen 1 deposition (Fig. 3B, C) that correlated with a significant increase of soluble collagen (Fig. 3D) and of its protein and mRNA levels (Fig. 3 E-G). Moreover, although the expression of fibronectin did not change significantly, an increment in the protein levels of SMA and in mRNA levels of Fn1, Acta2 and other fibrosis-related genes was observed in CDKO 3 days after UUO compared to WT mice (Fig. 3 F-H). The assessment of kidney function in WT and CDKO mice revealed a significant increase in BUN and no significant changes in blood creatinine levels in CDKO 3 days after UUO (Fig. 3 I). Analysis of changes in Cry1^+/-^; Cry2^+/-^ mice revealed a significant increase in the protein levels of Col1a1 (Fig. 3F), while mice deficient for each of the two Cry genes did not reveal statistically significant differences compared to WT mice after UUO (Fig. 3D-H). We take these results to propose that in the specific fibrosis model of UUO, the absence of Cry1/2 is a major determinant for phenotypic aggravation, while the latter is spared in mice genetically deficient for Bmal1 or Clock.

**Figure 3:**
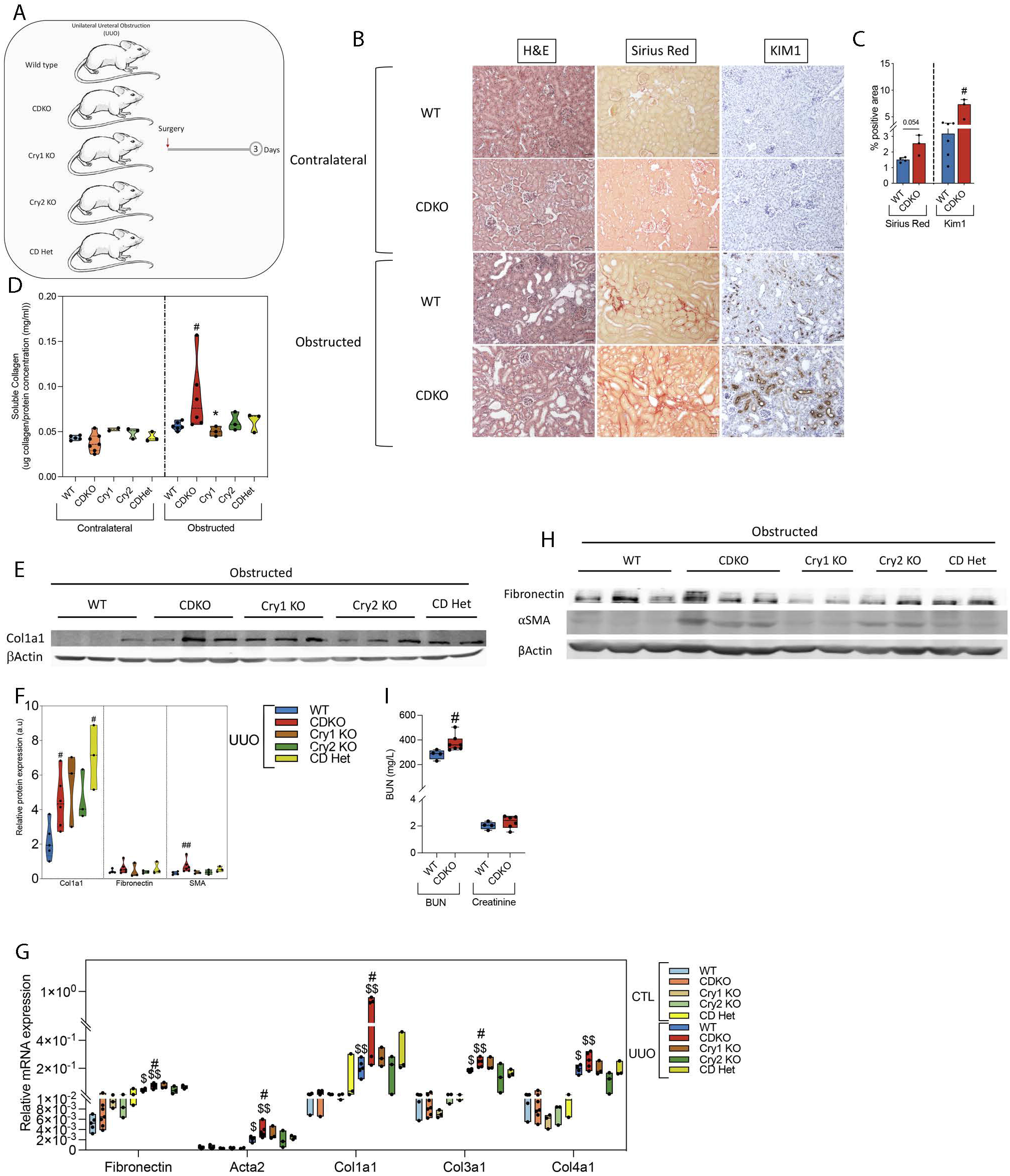
Fibrosis is exacerbated by Cry 1/2 genetic deficiency. (A) Schematic of experimental design of WT, CDKO, Cry1 KO, Cry2 KO and CD Het mice subjected to UUO for 3 days. (B) Representative microphotographs of H&E, Sirius red and KIM1 immunohistochemistry (IHC) stains from contralateral and obstructed kidneys of WT and CDKO mice 3 days after UUO. Scale bar: 100 μm. (C) Quantification of Sirius red and KIM1 IHC in kidney sections of the different genetically-modified mouse models denoted in (B), data are represented as the median with interquartile range. (D) Quantification of the total soluble collagen in frozen kidneys from WT, CDKO, Cry1 KO, Cry2 KO and CD Het mice subjected to UUO for 3 days, data are represented in a violin plot as the median with interquartile range. (E) Immunoblot depicting the expression of Col1a1 in kidneys from WT, CDKO, Cry1 KO, Cry2 KO and CD Het mice subjected to UUO for 3 days, β-actin levels were used for normalization. (F) Relative protein expression of Col1a1, Fibronectin and SMA of images in E and H, data are represented in a violin plot as the median with interquartile range. (G) Relative mRNA expression of fibrosis-related genes in WT, CDKO, Cry1 KO, Cry2 KO and CD Het mice subjected to UUO for 3 days, data are represented in a box plot as the median with interquartile range. (H) Immunoblot depicting the expression of αSMA and Fibronectin in kidneys from WT, CDKO, Cry1 KO, Cry2 KO and CD Het mice subjected to UUO for 3 days, β-actin levels were used for normalization. (I) BUN and Plasma creatinine levels of WT and CDKO mice 3 days after UUO, data are represented in a box plot as the median with interquartile range. Number of mice: Cry-deficient mice: WT (n=5), CDKO (n=6), Cry1 KO (n=3), Cry2 KO (n=3), CDhet mice (n=3); ^#^P<0.05, ^##^P<0.01 compared to the WT; ^*^P<0.05 compared to the CDKO; ^$^P<0.05, ^$$^P<0.01 compared to their respective contralateral kidney.

### Deletion of circadian components promotes differential patterns of inflammation in experimental models of kidney fibrosis

Fibrosis is the result of a maladaptive repair response to chronic inflammation. Thus, we set out to determine the role of the circadian clock in the inflammatory response that usually precedes and accompanies fibrosis. To do so, we evaluated the abundance of innate immune cells in the early (3 days UUO) and late (7 days UUO and 25 days ADN model) phases of inflammation using flow cytometry. Off note, the marker GR1 is comprised of two different membrane molecules, Ly6c and Ly6g. The first one is expressed either in monocytes and granulocytes, while Ly6g is specifically expressed in neutrophils^23^. Three days after UUO (Fig. 4A), we found that kidneys from Clock^Δ19^ mice showed a higher number of inflammatory macrophages (CD45+CD11b+F4/80+CD86+CD206-). Similarly, the monocyte population (CD45+CD11b+F4/80-Ly6c+) was also increased in these animals compared to WT mice, while no differences were observed in the neutrophil subset (CD45+CD11b+F4/80-CD86+Ly6c+Ly6g+) (Fig. 4 B-D). Consistently, markers of early inflammation (CD80, CD86, GM-CSF, iNOS) and levels of pro-inflammatory cytokines (TNF-α, IL1β, IL6, IFNγ) were increased in renal tissue of Clock^Δ19^ mice (Fig. 4 E, F).

**Figure 4:**
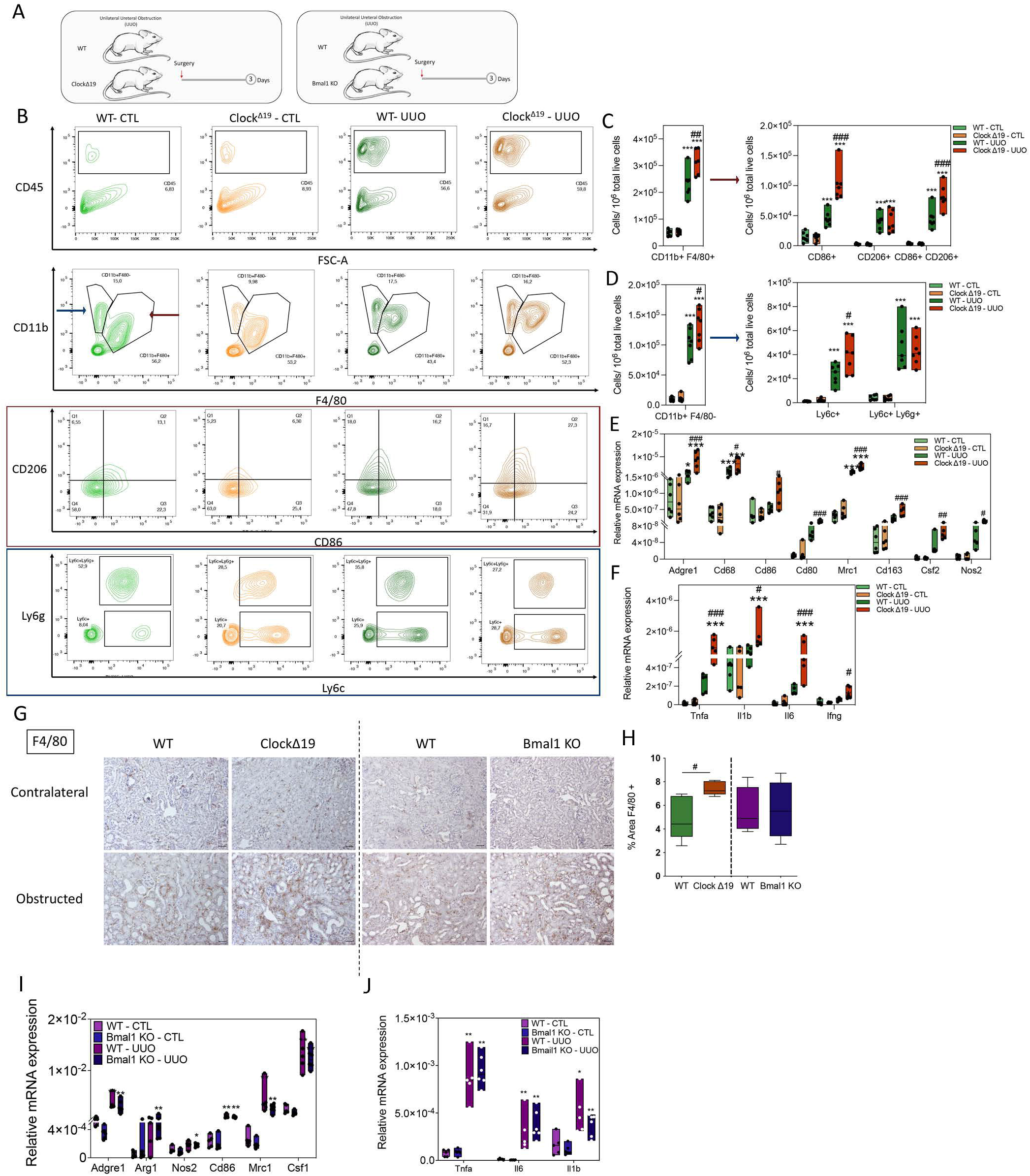
Clock^Δ19^ mice exhibit an increase in the number of CD86+ and CD86+/CD206+ macrophage and monocyte populations in the early phase of inflammation. (A) Schematic of experimental design of Clock^Δ19^ and Bmal1 KO mice subjected to UUO for 3 days. (B) Flow cytometry dot plots denoting the expression of inflammatory-related markers selected for the analysis of different innate immune populations. CD45 is a panleukocyte antigen also present in hematopoietic cells, monocytes and macrophages. F4/80 is a surface marker expressed in monocytes and macrophages and CD11b is a marker commonly expressed in inflammatory myeloid cells. CD86 and CD206 were used for the selection of proinflammatory or antiinflammatory macrophage subpopulations (CD45+F4/80+CD11b+), respectively. Ly6c and Ly6g were used for the selection of monocytes (CD45+F4/80-CD11b+Ly6c+Ly6g-) and neutrophils (CD45+F4/80-CD11b+Ly6c+Ly6g+) in kidneys from Wt and Clock^Δ19^ mice 3 days after UUO. (C, D) Quantification of the different immune populations analyzed in (B), data are represented in a box plot as the median with interquartile range. (E, F) Relative mRNA expression of inflammatory-related genes in kidneys from Wt and Clock^Δ19^ mice 3 days after UUO, data are represented in a box plot as the median with interquartile range. (G) Representative pictures of F4/80 immunohistochemistry stain (IHC) of contralateral and obstructed kidneys of Bmal1 KO, Clock^Δ19^ and their corresponding WT mice, 3 days after UUO. Scale bar: 100 μm. (H) Quantification of F4/80 IHC in obstructed kidney sections of the different mouse models denoted in (F); data are represented in box plots as the median with interquartile range. (I, J) Relative mRNA expression of inflammatory-related genes in kidneys from Wt and Bmal1 KO mice 3 days after UUO, data are represented in a box plot as the median with interquartile range. Number of mice: Clock deficient mice: WT (6) and Clock^Δ19^ mice (6); Bmal1 deficient mice: WT (5) and Bmal1 KO (6). ^*^P<0.05, ^**^P<0.01 ^***^P<0.001 compared to their respective contralateral kidney; ^#^P<0.05, ^##^P<0.01, ^###^P<0.001 compared to WT.

Immunohistochemical evaluation of the macrophage population confirmed their increased presence (Fig. 4 G, H). The CD86 and CD206 double positive subset of macrophages - a transitional phenotype from a pro-inflammatory to antiinflammatory state-was also increased in these mutant mice compared to WT (Fig 4 B, C, E). The number of CD86-CD206+ anti-inflammatory macrophages remained invariable between genotypes (Fig. 4 B, C). Similarly, Th2 antiinflammatory cytokines (IL-4, IL-10) were undetectable (data not shown). In contrast, in mice with a global deletion of Bmal1 we did not observe an increase in inflammation 3 days after UUO (Fig. 4 G-J).

Studies in the late phase of inflammation were also done in WT mice (normal phenotype) and Clock^Δ19^ mice (Fig S8A, E). We observed that in the ADN model, the alteration of Clock was associated with increased presence of antiinflammatory CD86-CD206+ macrophages as well as of CD86 and CD206 double positives. On the contrary, no significant differences were observed in the proinflammatory CD86+CD206-macrophages and CD45+CD11b+F4/80-Ly6c+ monocytes (Fig. S8 B-D). Consistently, we found increased abundance of inflammatory markers (Arg1, Mrc1, IL-4, IL-17) and of the profibrogenic cytokine TGF-□ in kidneys from Clock^Δ19^ mice compared to those of WT mice 7 days after UUO (Fig. S8 F, G). We then explored the contribution of Bmal1 in a mouse model with conditional suppression of the Bmal1 gene in the renal tubule (Bmal1 cKO) (Fig S8H). We did not observe significant differences in the abundance of macrophage markers in mice subjected to UUO (Fig. S8I). We found elevation of IL-10, IL-13 and IL-17 in Bmal1 cKO mice, although only IL-13 reached statistical significance (Fig. S8J). Mice with a global deletion of Bmal1 (Bmal1 KO) (Fig. S8K) did not show significant differences regarding the presence of macrophages or the levels of TNF-□ (Fig. S8L). These data further support an involvement of Clock, rather than Bmal1, in the early and late phase of macrophage inflammatory response inherent to the UUO model.

To find out if the absence of Cry1, Cry2 or both did not only aggravate fibrosis but was also related to the accompanying inflammatory response, we studied the population of inflammatory cells in CDKO mice, 3 days after UUO (Fig. 5A). We found a significant increase in the innate immunity related CD11b+F4/80-GR1+ inflammatory cell population in obstructed kidneys of CDKO mice (Fig. 5 B, C). This was not reflected in an increase of the macrophage populations in the same condition (Fig. 5 D). The analysis of mRNA levels of Ly6c and Ly6g markers revealed that CDKO mice exhibited a significant increase in both either in control and obstructed kidneys, a phenomenon not present in animals deficient for only one of the Cry genes or in the Cry1 and Cry2 double heterozygous mice (Fig. 5 E). The presence of several macrophage-related markers was increased in obstructed kidneys in all the genotypes analyzed (Fig. 5F), thus suggesting no differential effect related to the absence of Cry at this stage of inflammation. Overall, these data suggest that the absence of Cry1/Cry2 selectively affects the abundance of GR1+ cells, irrespective of kidney damage, in the early inflammatory response accompanying UUO.

**Figure 5:**
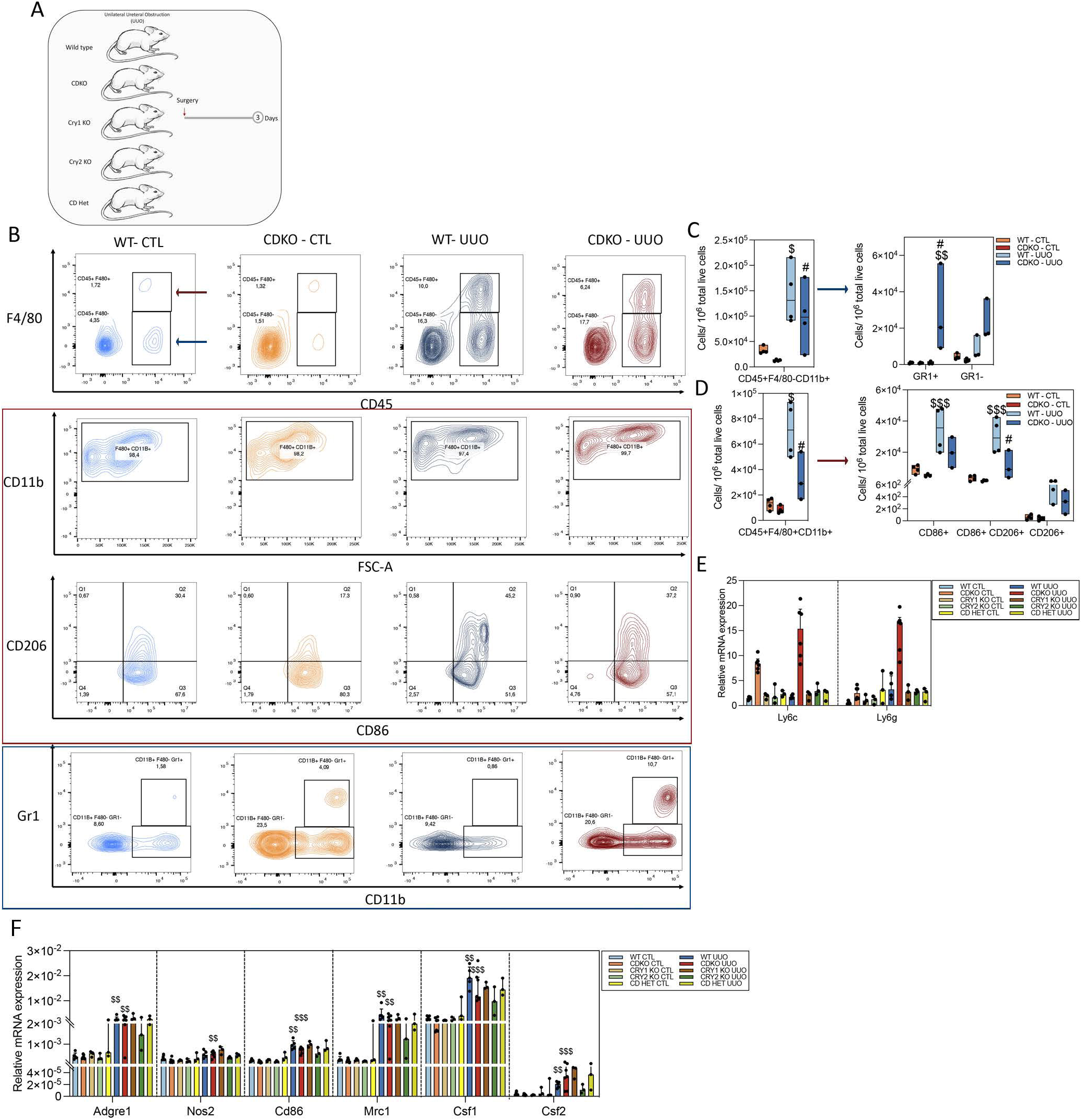
The inflammatory response of CDKO mice is associated with an increased presence of neutrophils. (A) Schematic of experimental design of WT, CDKO, Cry1 KO, Cry2 KO and CD Het mice subjected to UUO for 3 days. (B) Flow cytometry dot plots denoting the expression of inflammatory-related markers selected for the analysis of different innate immune populations (see legend of Figure 4 for detailed explanation).Gr1 was used for the selection of either monocytes and neutrophils (CD45+F4/80-CD11b+Gr1+) in kidneys from WT (n=3) and CDKO (n=3) mice, 3 days after UUO. (C, D) Quantification of the different immune populations analyzed in (B), data are represented in a box plot as the median with interquartile range. (E, F) Relative mRNA expression of myeloid inflammatory-related genes in kidneys from WT, CDKO, Cry1 KO, Cry2 KO and CD Het mice subjected to UUO for 3 days, data are represented in a box plot as the median with interquartile range. Number of mice: WT (n=5), CDKO (n=6), Cry1 (n=3), Cry2 (n=3) and CDhet (n=3) mice. ^#^P<0.05, compared to the WT; ^$^P<0.05, ^$$^P<0.01, ^$$$^P<0.001 compared to their respective contralateral kidney.

### The expression of metabolism-related genes is differentially affected by the selective deletion of molecular clock components in the UUO model of kidney fibrosis

In the past years the role of metabolic failure of tubular epithelial cells in the genesis and progression of kidney disease has been clearly established ^6,7^. Due to the connection between CR and metabolism, we sought to determine the effect of the deletion of clock components on the expression of key metabolic genes in the UUO model of kidney fibrosis. Specifically, we analyzed genes involved in five major metabolic pathways, including those related to mitochondrial function and bioenergetics. We found that CDKO (Fig. 6A) showed a significant reduction in transcripts of all these five metabolic routes both in non-obstructed and obstructed kidneys (Fig. 6B). Consistent with this transcriptomic profiling, TFAM protein levels were diminished in contralateral and obstructed kidneys of the CDKO mice, compared to WT and single Cry1 or Cry2 null mice, while no significant differences were observed in Cpt1a protein expression (Fig. 6 C-E). Interestingly, the mitochondrial DNA copy number was increased in CDKO kidneys in both, control and damaged kidneys (Fig. 6 F). By contrast, the number of genes with significant reduction in their expression was much lower in the case of the Bmal1 KO mice, mostly affecting FAO-related genes (Fig. S9A-D). This pattern was not maintained in the Clock^Δ19^ mice (Fig. S9E-H) or Bmal1 cKO (Fig. S9I-L), thus suggesting that the clock regulators Cry1/2 are specifically involved in the metabolic derangement associated to kidney damage.

**Figure 6:**
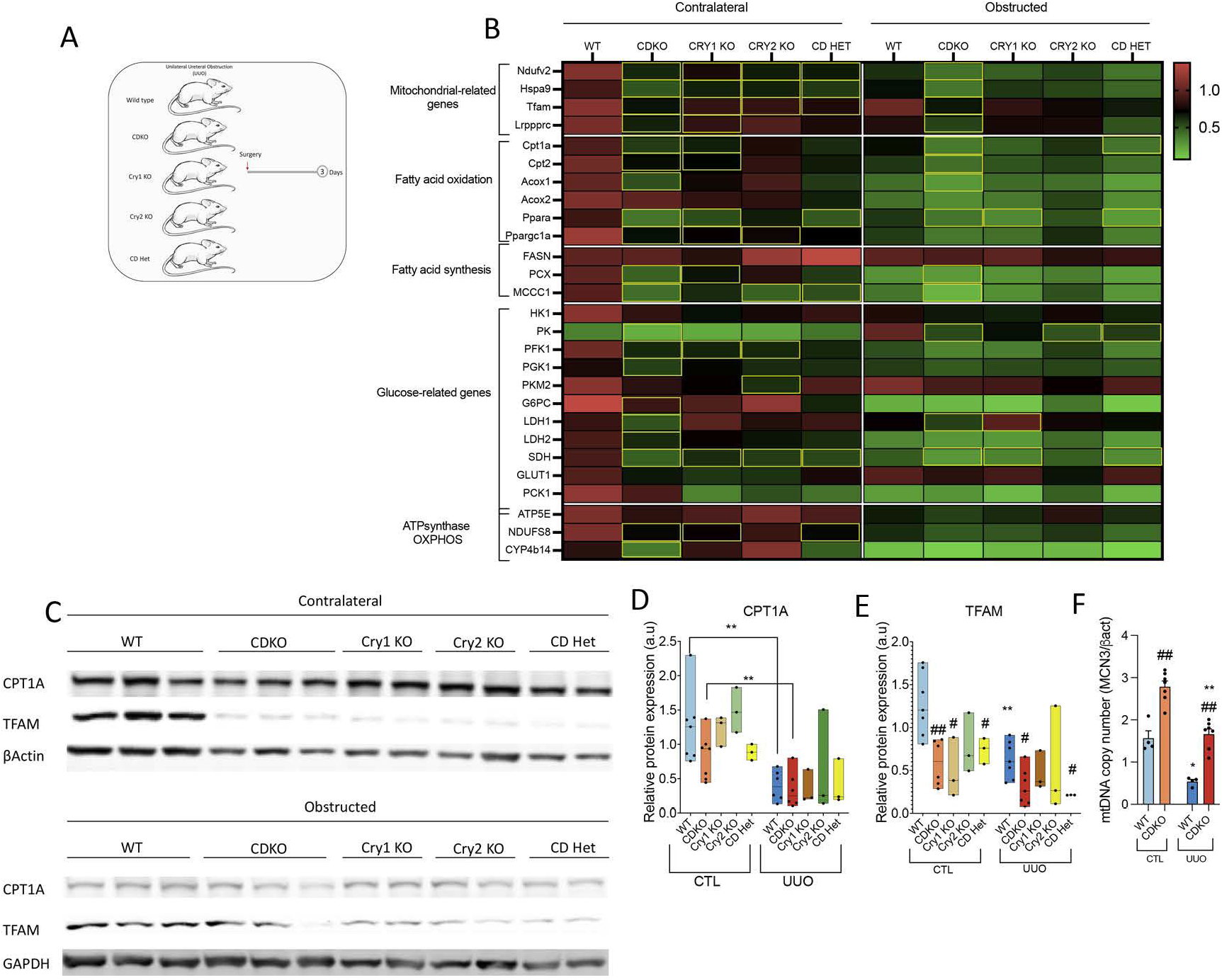
The expression profile of metabolism and mitochondrial-related genes is significantly altered in CDKO mice. (A) Schematic of experimental design of WT, CDKO, Cry1 KO, Cry2 KO and CD Het mice subjected to UUO for 3 days. (B) Heat map of normalized expressions of metabolism and mitochondrial-related genes in CDKO mice, in kidneys 3 days after UUO. Yellow rectangles denote significant differences compared with kidneys from Wt mice. (C) Immunoblot depicting the expression of Cpt1a and Tfam proteins in kidneys from mice with the different genotypes of Cry1 and Cry2 absence, 3 days after UUO; β-actin and Gapdh levels were used for normalization. (D, E) Relative protein expression of (D) Cpt1a and (E) Tfam from the immunoblots showed in (B). Data are represented in a box plot as the median with interquartile range. (F) Relative mitochondrial DNA copy number was analyzed in kidneys from Wt and CDKO 3 days after UUO. Data are represented as the median with interquartile range. Number of mice: WT (n=5), CDKO (n=6), Cry1 KO (n=3), Cry2 KO (n=3). ^*^P<0.05, ^**^P<0.01 compared to their respective contralateral kidney; ^#^P<0.05, ^##^P<0.01 compared to the Wt.

## Discussion

The progression of CKD is associated with sleep disturbances and an irregular circadian pattern of hormone secretion and blood pressure oscillation^12,24–27^. However, the mechanisms underlying these alterations remain obscure and plausibly, an altered biunivocal relationship between kidney function and circadian rhythm lies at the core of this dysfunction. Our results demonstrate a disruption of the peripheral molecular clock in the kidneys as a consequence of renal damage. Interestingly, we observed that Bmal1 expression was steadily upregulated in damaged kidneys from the three different models analyzed. Moreover, we found that TGFβ1 increased the expression of core clock genes in HPTEC cells in a time-dependent fashion, with a marked effect on Bmal1. The prevention of this effect by the pharmacological inhibitor of the Alk 4-5-7 TGFβ receptor SB505124 supports the involvement of this pathway, strongly suggesting the role of Alk 5, given its higher affinity for the inhibitor compared to ALK4 and 7^28^. Our results are consistent with other reports that found a perturbed pattern in the rhythmic oscillations of circadian genes in injured kidneys^11,29–31^. In addition, previous studies have revealed that the TGFβ pathway is able to modify the oscillatory patterns of core clock genes from different tissues^32,33^ and other authors have reported a reduction in the expression of Bmal1 in heart tissue of Smad3 knockout mice^34^ as well as a role for Smad4 in pancreatic cancer cells^35^. However, the specific mechanism has not yet been clarified. Although we did not find increased transcriptional activity of several potential cis-regulatory elements after Smad3 overexpression, we cannot exclude the participation of other as yet, unidentified enhancers. Hence, further studies are needed to decipher the molecular mechanisms responsible for Bmal1 upregulation in kidney fibrosis.

Our results revealed a worsening of fibrosis in the CDKO mice, while less collagen deposition was found in the Bmal1 KO and no differences were appreciated in the tubule-specific Bmal1 KO and Clock^Δ19^ mice. In the literature, variable and controversial results have been reported regarding the role of Bmal1 and Clock in the instauration of fibrosis. While a pro-fibrotic role of these proteins was observed in some studies in the context of kidney, lung and heart disease^29,33,36^, Bmal1 and Clock appear to exert a protective role in other studies related to kidney, lung and liver fibrosis^11,37–40^. In contrast, our observations in the CDKO mice are concordant with studies in other fibrotic contexts. Hence, mice with a deletion of either Cry1 or Cry2 developed a more severe phenotype in a barium chloride-induced model of muscle fibrosis^41^ and Cry1-null, Cry2-null mice have shown altered expression patterns of aldosterone, a vital extrarenal hormonal regulator of kidney-mediated hydroelectrolytic homeostasis^42^.

A persistent inflammatory response leads to the perpetuation of repair processes that involve a gradual instauration of fibrosis^43^. While no substantial differences were observed in our study regarding Bmal1, our data revealed a relevance of Clock and Cry in the macrophage and neutrophil populations, respectively. To our knowledge, this is one of the first studies documenting the relevance of these core clock genes in kidney inflammation. In other pathological contexts, Bmal1 seems to play a role in the metabolic switch of macrophage populations^44–46^. However, Bmal1 was not critical for the inflammatory instauration associated to lung and liver disease^38,47^. In contrast, Clock^Δ19^ mutation has been associated with a pro-inflammatory phenotype in non-alcoholic liver disease^40^. Clock has been identified as a protein implicated in the NF-□-dependent inflammatory response by its direct interaction with the NF-□ subunit, p65, which competes with Bmal1^48^. These authors observed that Clock KO mice showed a decrease in the accumulation of NF-□ in hepatocyte nuclei, a phenomenon absent in the Clock^Δ19^ genotype, suggesting a Clock-mediated circadian-independent regulatory mechanism of inflammation. Although these studies are consistent with our observations, the absence of significant changes in the fibrotic phenotype observed by us and the variability in the results reported by other authors^11,37–40^ could imply that Clock-mediated inflammatory response is unable by itself to promote a drastic fibrotic phenotype. Neutrophils are subjected to circadian regulation as illustrated by diurnal variations in their levels^50^. However, the relevance of the different core clock components in this regulation is poorly understood. To our knowledge, a specific role for Cry proteins in the regulation of neutrophil-mediated inflammatory responses has not been previously reported. Of interest, the neutrophilic presence beyond the acute phase of inflammation has been linked to enhanced renal fibrosis^52,53^. In keeping, in consistence with our observations, a massive infiltration of leukocytes into the lungs and kidneys has been previously reported in CDKO mice as a consequence of an autoimmune response through the modulation of the BCR-signaling pathway^51^. Whether this dysfunctional B cell response contributes to the observed aggravation of kidney fibrosis remains to be investigated. Moreover, Cry proteins have also been associated with the inhibition of the NF-κB-mediated inflammatory response^49^.

Alterations of mitochondrial function and cell bioenergetics have been linked to the development of several diseases^54–57^. Our results showed an overall reduction in the expression of most of the metabolism-related genes analyzed in the CDKO mice and a decreased expression of some FAO and glucose-related genes in the Bmal1 KO. In consistence with our data, the absence of Bmal1 has been associated with metabolic alterations in non-damaged kidneys^17^, as well as to pathological alterations in other tissues^37,38^. Furthermore, our data revealed that Tfam levels were dramatically reduced in CDKO mice. Tubule-specific deletion of Tfam has been related to an increase in renal fibrosis^58^, thus supporting the pro-fibrotic phenotype found in these mice. While these results support a link between Cry function and kidney metabolism, the role of Cry proteins in the metabolic derangement associated with fibrosis should be further explored. Taken together, our results sustain that kidney damage promotes a profound alteration in the expression of the circadian clock components and suggest the existence of a direct regulation of the core clock by TGFβ1. In addition, they support a major protective role for Cry proteins in renal fibrosis, together with a role of both Cry and Clock in the accompanying neutrophil and macrophage inflammatory response, respectively (Graphical abstract).

## Acknowledgements

This work was supported by grants from the Ministerio de Ciencia e Innovación PID2019-104233RB-100/AEI/10.13039/501100011033 (SL), Instituto de Salud Carlos III REDinREN RD12/0021/0009 and RD16/0009/0016 (SL), Comunidad de Madrid “NOVELREN” B2017/BMD-3751 (SL, CB) and Fundación Renal “Iñigo Alvarez de Toledo” (SL), all from Spain. Carlos Rey-Serra is the recipient of an FPI research training contract from the Spanish Research State Agency (BES-2016-076735). The CBMSO receives institutional support from Fundación “Ramón Areces”. We thank the laboratories of Fernando Rodríguez Pascual (CBMSO) for help with plasmid constructions and of Marta Ruiz Ortega at the Fundación Jiménez Díaz for help with immunohistochemistry. We also thank the help of the following facilities of the CBMSO: animal housing, flow cytometry and confocal and electron microscopy.

---

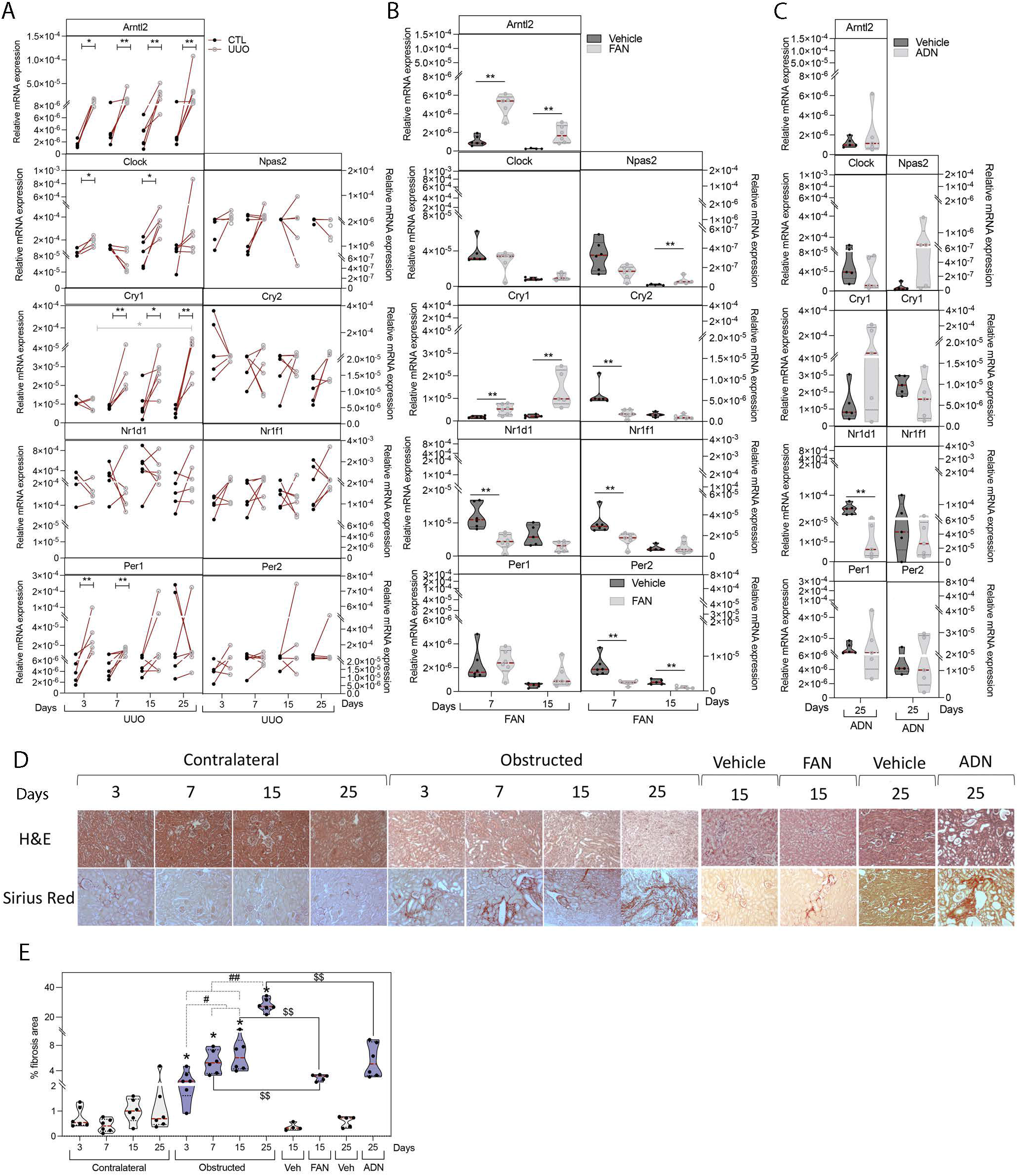

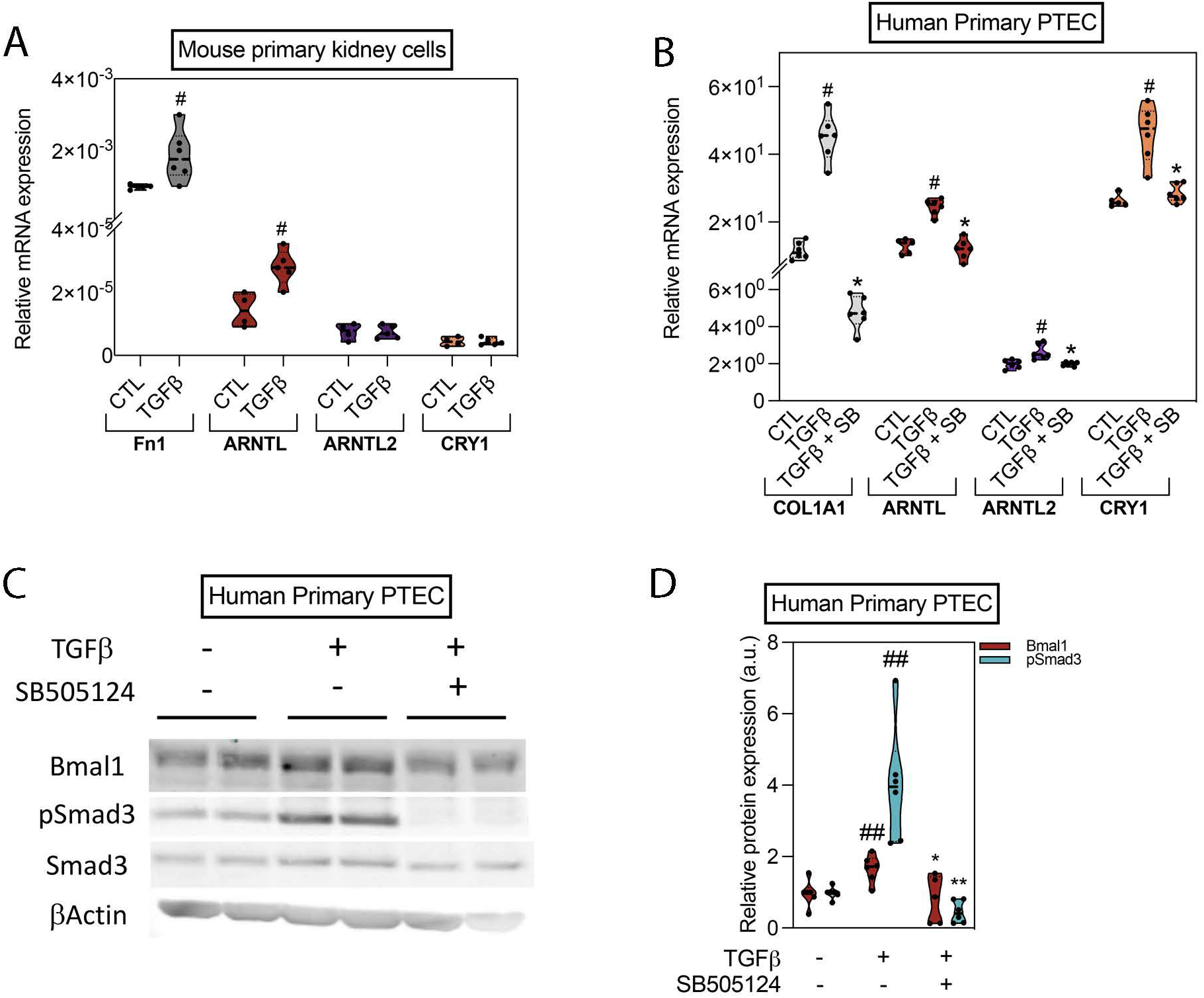

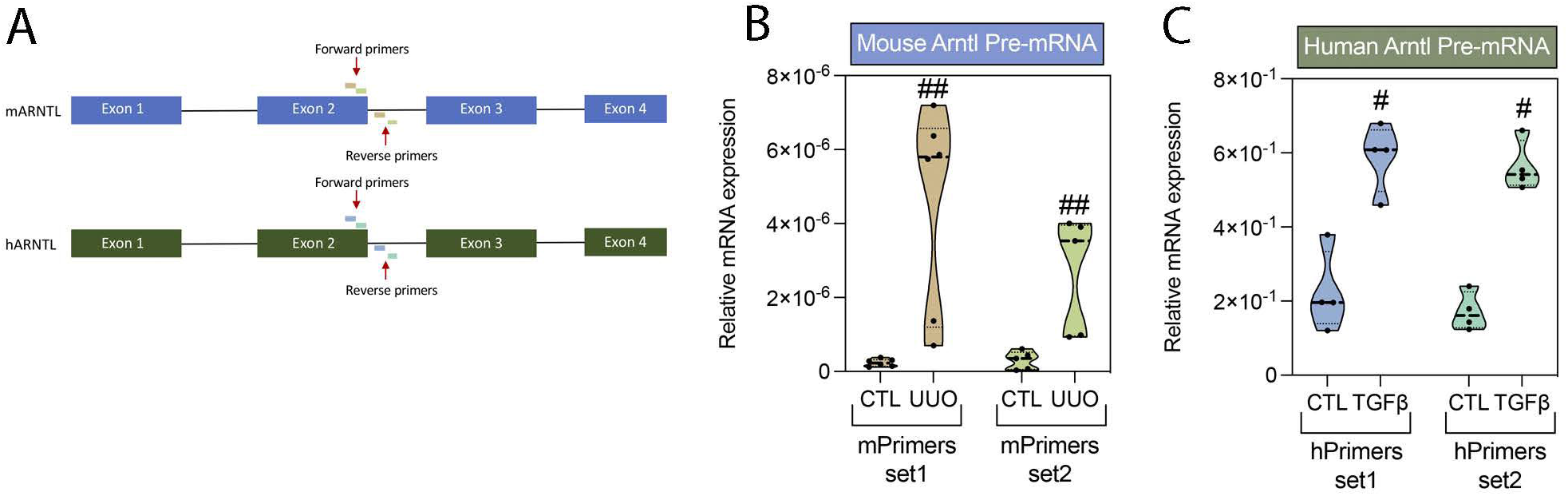

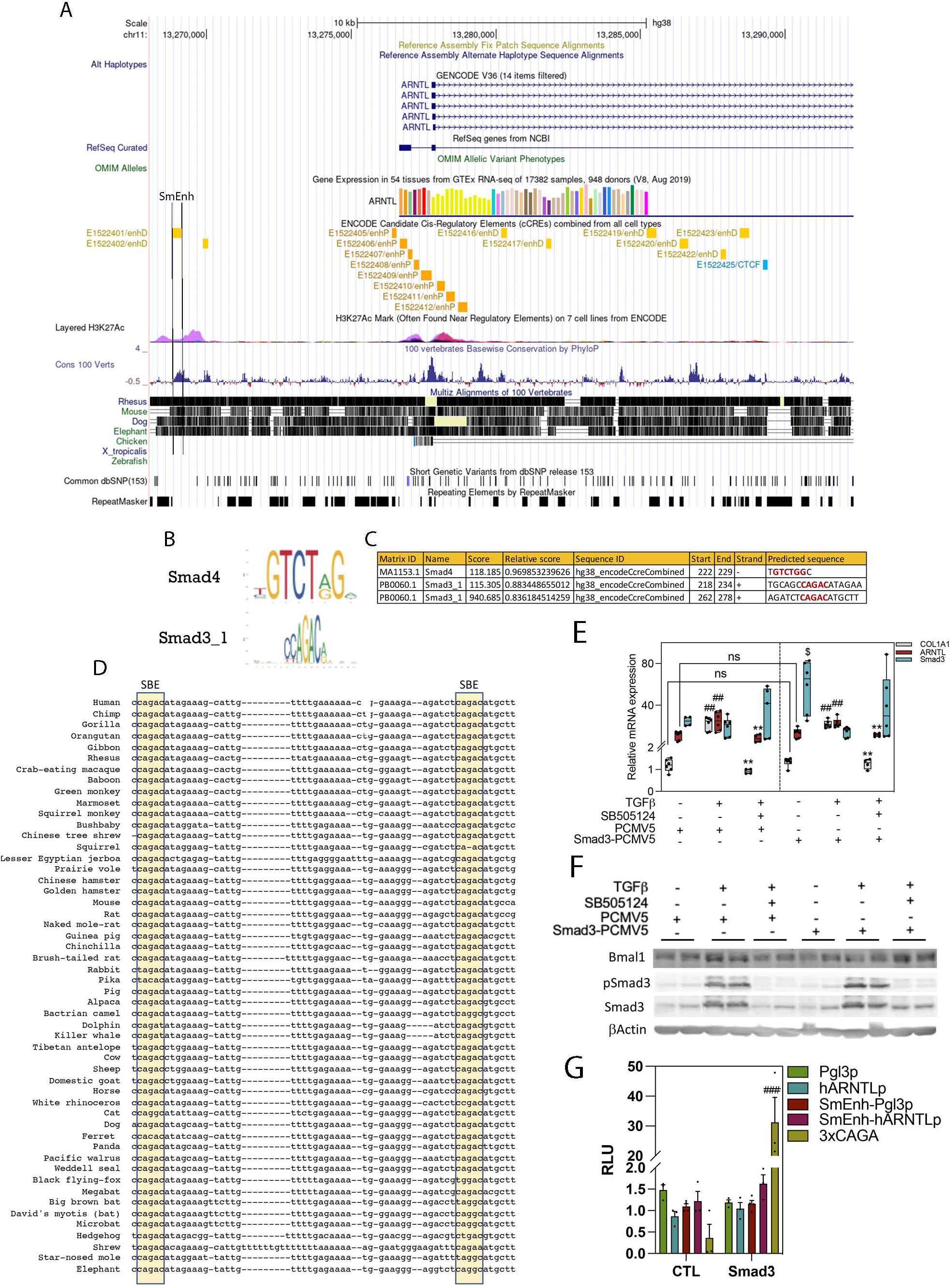

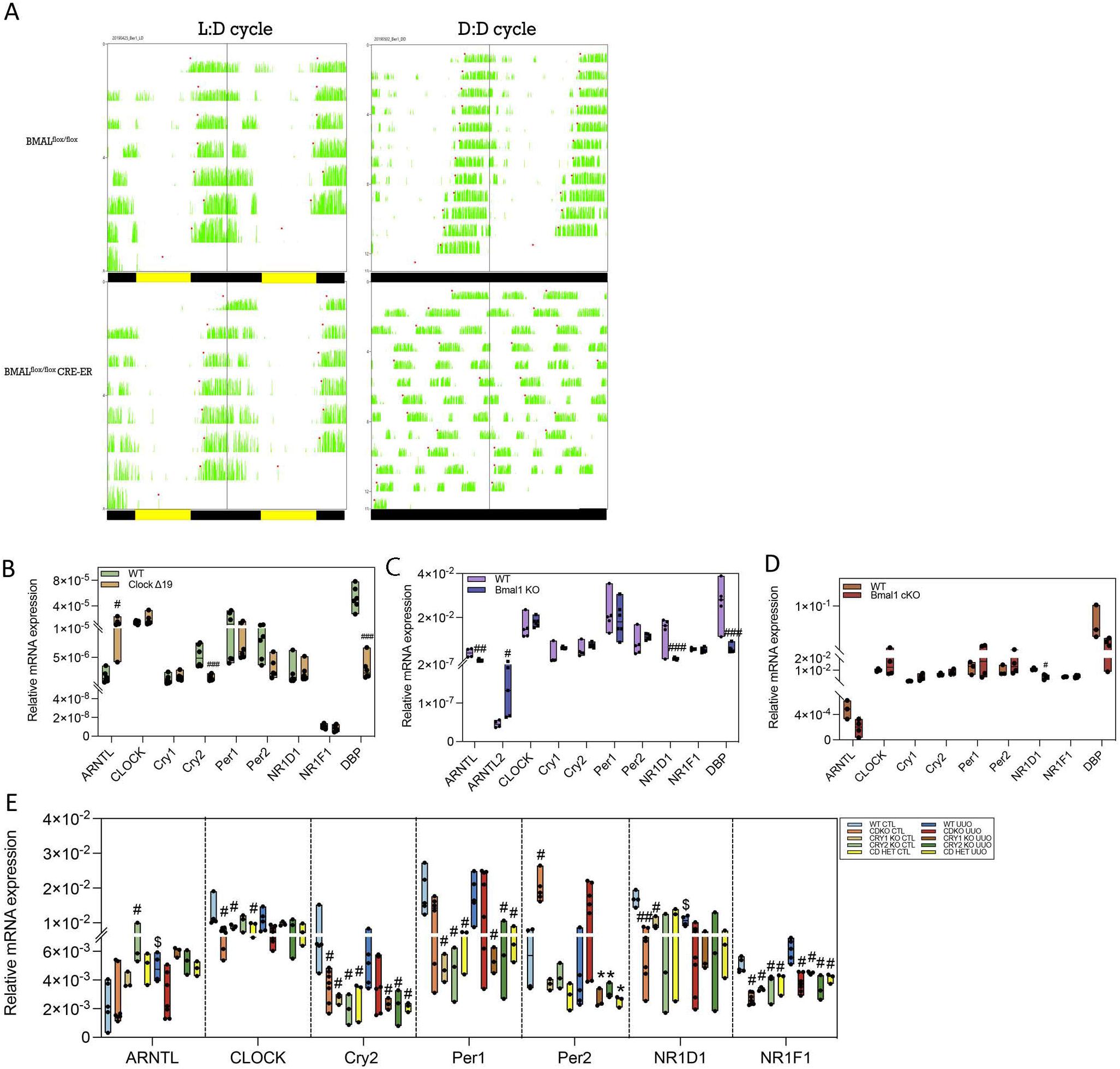

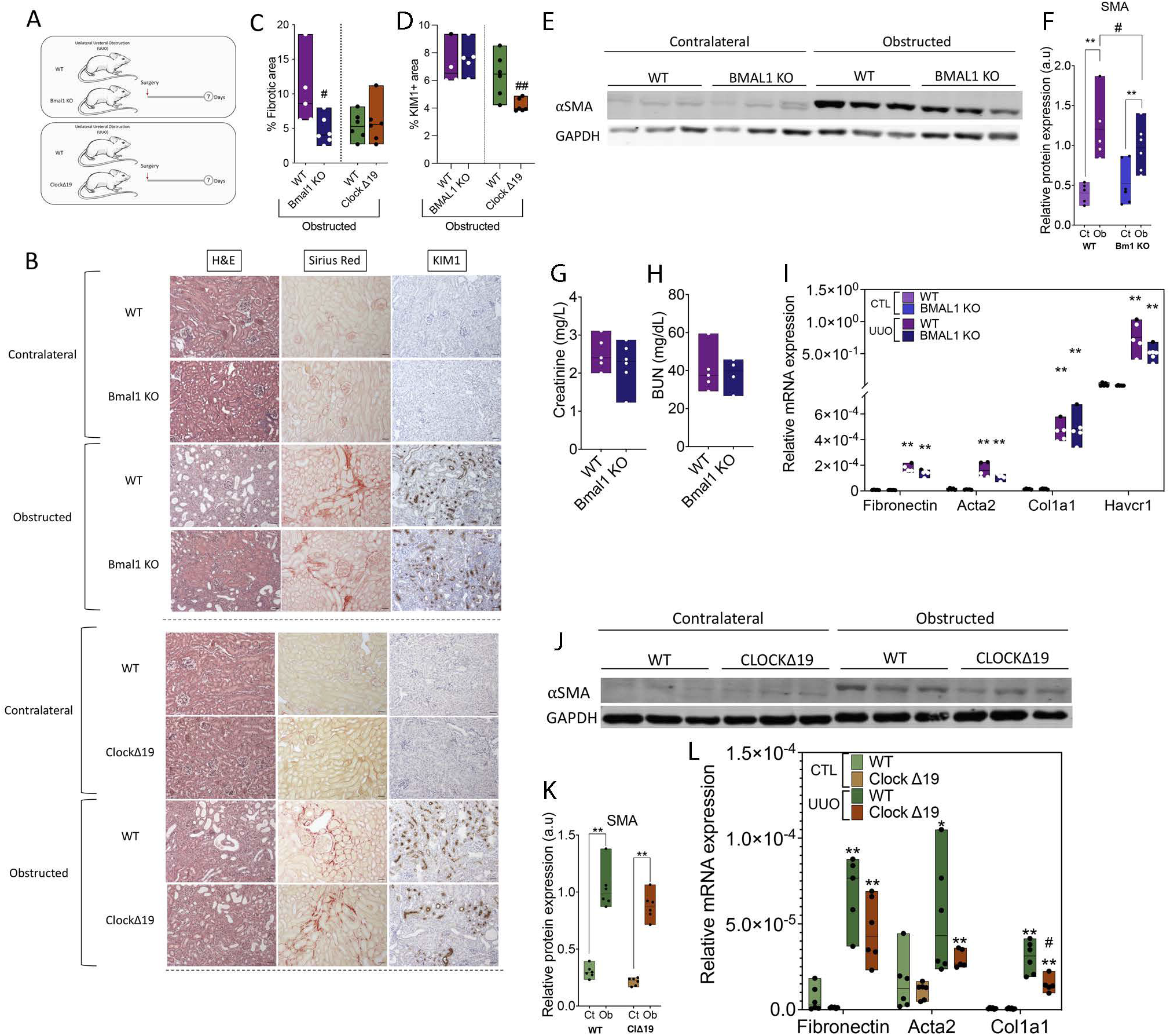

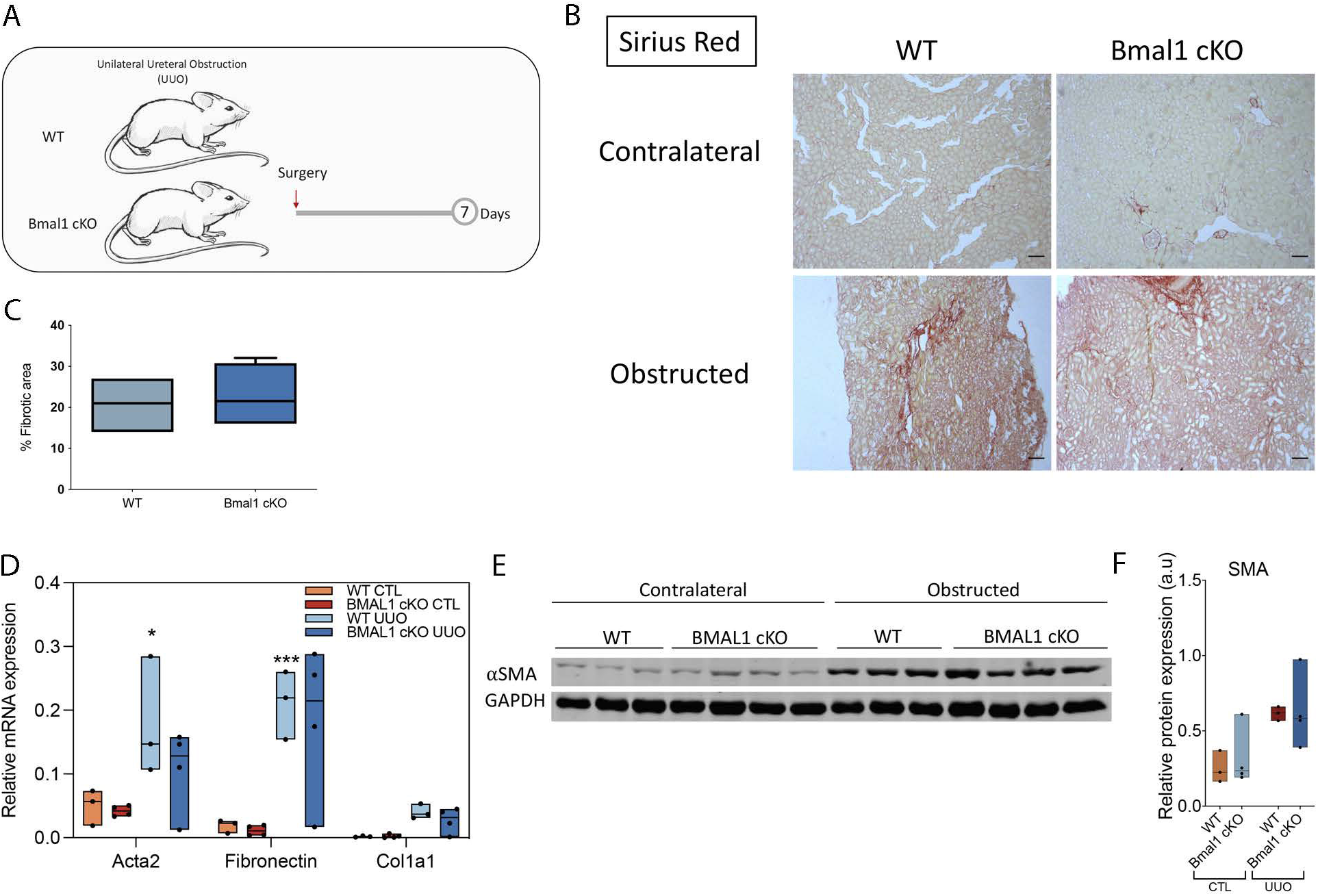

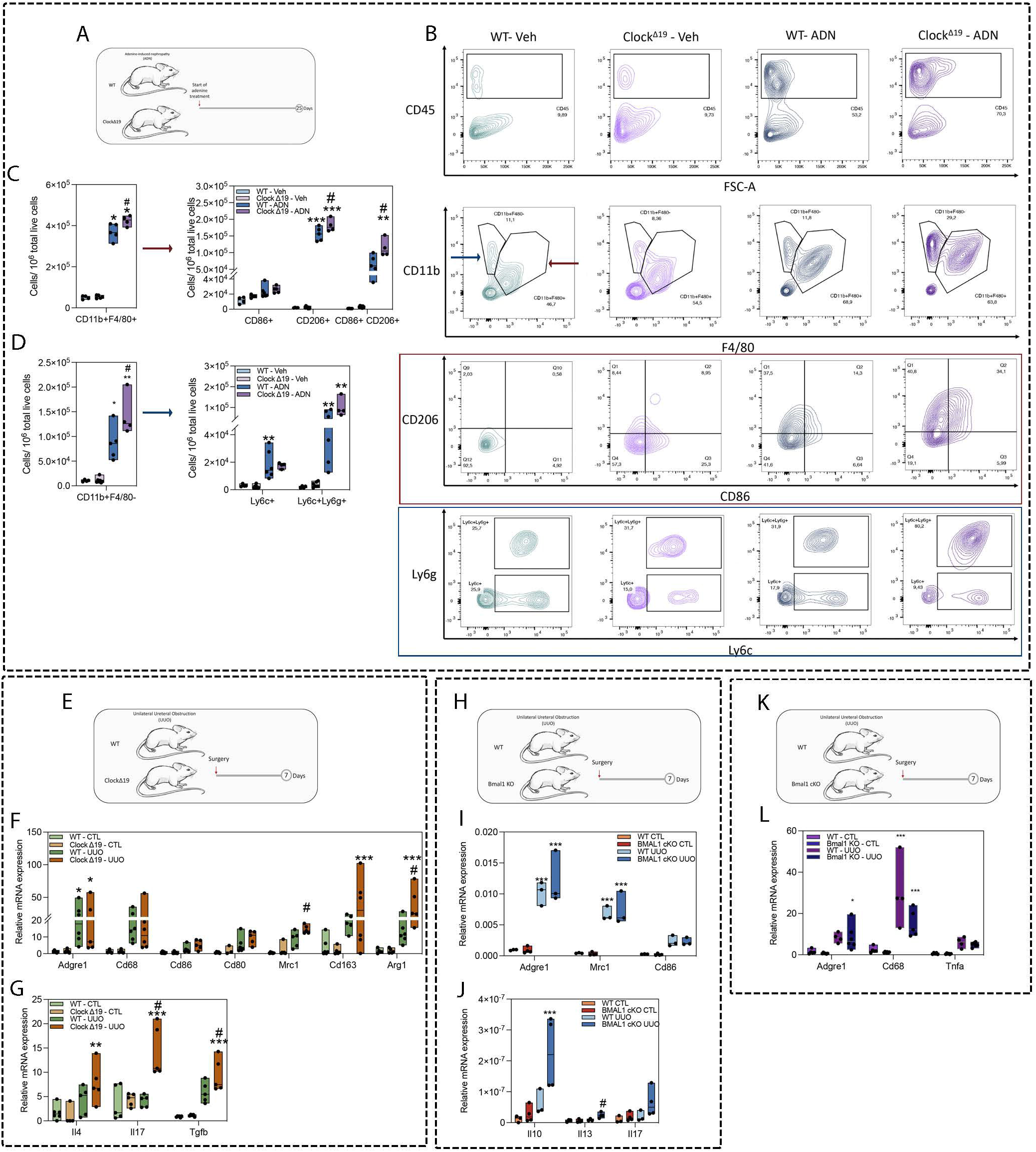

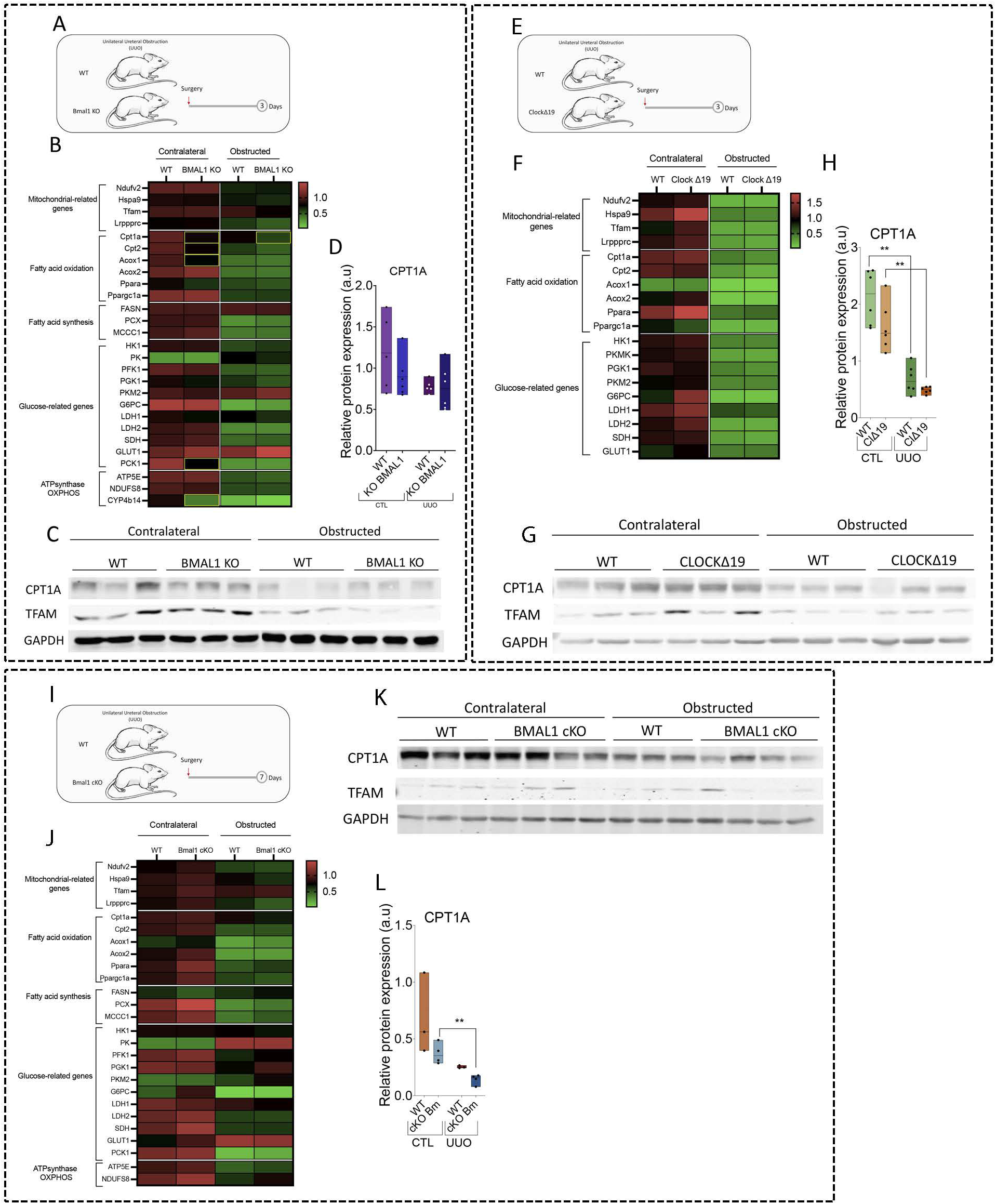

