## Supplemental methods and Tables for "Cross regulation between the molecular clock and kidney inflammatory, metabolic and fibrotic responses"

**Supplementary methods**

**Ethical compliance for mouse handling and experimental models.** Mice were housed in the Specific-pathogen-free (SPF) animal facility at the CBMSO or Salk Institute in accordance with the EU and US regulations for all the procedures. Animals were handled in agreement with the Guide for the Care and Use of Laboratory Animals contained in Directive 2010/63/EU of the European Parliament and in the Animal Welfare Act of the United States Department of Agriculture. Approval was granted by the local ethics review board of the Centro de Biología Molecular "Severo Ochoa" in Madrid, the Ethics committee at CSIC and the Regulatory Unit for Animal Experimental Procedures from the Comunidad de Madrid and by the Institutional Animal Care and Use Committee (IACUC) of the Salk Institute.

**Assessment of kidney function.** Blood urea nitrogen (BUN) was analyzed in serum samples by using the BUN Colorimetric Detection Kit. Serum creatinine was measured by capillary electrophoresis (Agilent 7100) coupled to TOF mass spectrometry (Agilent 6224) [56,57]. A calibration curve was used for quantification with creatinine; methionine sulphone was the internal standard.

**Animal Handling**

The genotype of the animals was confirmed by PCR by using the primers listed in **Table S1**. The animals were maintained *ad libitum* on regular chow diet, under constant temperature and humidity in a 12-hour light/dark cycle. Doxycycline and Tamoxifen were used for the induction of Cre expression in 8 weeks-old Bmal1 cKO and Bmal1 KO mice, respectively. Mice were treated daily with 2 mg/ml of doxycycline in drinking water [17] or 3.7 mg of tamoxifen administered by oral gavage for 2 weeks or 5 days, respectively. Subsequent experiments were performed four or two weeks after the end of the treatment, respectively, in order to avoid potential side effects related to their administration.

**Analysis of the circadian activity disruption in the Bmal1 KO mice**

Mouse activity was tested in the Bmal1 KO and their WT mice after Cre expression induction with tamoxifen using the Mouse Home Cage Running Wheel (The Columbus Instruments, OH, USA).

Wheels revolutions were recorded by a computer through a sensor connected to the wheels. The mice were previously adapted to the running wheel cages for two days in a 12h light:12h dark cycle prior the analysis. Then, mouse activity was measured for two days in the same conditions and another 2 days in constant darkness.

**Mouse models of chronic kidney disease.** Unilateral Ureteral Obstruction (UUO) procedure was performed as described previously[58]. Briefly, 12-14 weeks old mice were anesthetized with 2% isoflurane. The hair in the abdominal area was shaved. An incision was made in the abdominal wall to expose the left kidney, the ureter was ligated twice and severed between the two ligatures and the kidney was returned gently to its place. Finally, the abdominal incision was closed with sutures and buprenorphine was used as an analgesic. Folic acid nephropathy (FAN) was carried out as previously described[59]. Mice were given 250 mg/kg of folic acid intraperitoneally (i.p) (Sigma-Aldrich) resuspended in 0.3 M sodium bicarbonate (vehicle) and control mice were given 0.1 ml of the vehicle. For adenine- induced renal failure (ADN), mice were given 50 mg/kg of adenine in 0.5% carboxymethyl cellulose (CMC) (Wako Pure Chemical Industries Ltd., Osaka, Japan) daily by oral gavage as described previously[60]. Control mice were given 0.1 ml of 0.5% CMC (vehicle). Mice were sacrificed 3, 7, 15 or 25 days after UUO; 7 and 15 days after FAN or 25 days after ADN. The three procedures and the sacrifice of the mice were performed at ZT9 (where ZT0 means light on and ZT12 means light off). Blood samples were collected right after the sacrifice by cardiac puncture and kidneys were collected after perfusion with PBS. Number of mice described in **Table S2**.

**Histological and immunohistochemical analysis.** Kidney samples were fixed in 4% neutral buffered formalin before being embedded in paraffin. H&E and Sirius red were performed on 5 µm sections using standard procedures. Sections of 3 µm were deparaffinized for IHC and antigen retrieval was performed with 10 mM citrate sodium buffer by using the PT Link (Dako, Santa Clara, CA, USA). Endogenous peroxidase and non-specific protein binding sites were blocked with 3% $H_2O_2$ and 4% bovine serum albumin (BSA) in 1X EnVision wash buffer (Dako, Santa Clara, CA, USA), respectively. Incubation with the following primary antibodies: KIM1 (AF1817, R&D systems, Minnesota, USA) and F4/80 (F4/80 (D2S9R) XP® Rabbit mAb #70076, Cell Signaling,

Massachusetts, USA) was carried out overnight at 4ºC and, right afterwards, incubation with biotinylated goat anti-mouse or anti-rabbit IgG was performed for 1h at 4ºC. VECTASTAIN ABC Kit (Vector Laboratories, Burlingame, CA, USA) was used for detection of the biotinylated-secondary antibodies. Tissue sections were revealed with 20 µg/ml 3,3'-diaminobenzidine (DAB, Dako) and counterstained with hematoxylin. Images were taken at 10x magnification with a Nikon's Eclipse TE2000-U light microscope (Nikon Instrument Europe B.V, Badhoevedorp, The Netherlands). The intensities of Sirius red (collagen deposition) and IHC were quantified automatically with the Image-pro Plus software (Media Cybernetics). Data are represented as the percentage of the ratio of positive area to total tissue.

**Plasmid construction**

*In-silico* analysis was performed on human ARNTL regulatory region using the R package for JASPAR2018[19]. The distal enhancer E1522401 (SmEnh) of the human ARNTL gene (located from -9325 to -8376; where +1 bp is the putative transcription start site (TSS)) was cloned into the Pgl3-promoter or Pgl3-basic vector (Promega Corporation, Madison, WI, USA). The human Arntl promoter region (hArntlp; from -1220 to +8) was used as the promoter sequence for the Pgl3-basic vector by cloning it downstream of the SmEnh sequence. For that purpose, human genomic DNA was obtained from HPTEC cells using the PureLink kit (Invitrogen, Carlsbad, CA, USA) according to manufacturers' instructions. The inserts were amplified by PCR using the primers listed in **Table S3** and the ACCUSTART kit (QuantaBio, VWR International, Pennsylvania, USA) following the manufacturers' instruction. The PCR products were tested by 1% agarose gel electrophoresis (CONDA, Madrid, Spain) and purified with QIAquick Gel Extraction Kit (Qiagen, Hilden, Germany). Once purified, the inserts were first cloned into PCR2.1 vector using the TA-Cloning kit (Invitrogen, Carlsbad, CA, USA) and subsequently subcloned into the PGL3-basic or PGL3-promoter vectors. The construct was verified by Sanger sequencing and diagnostic enzymatic digestions with restriction enzymes.

**Cell culture.** Immortalized renal human proximal tubule epithelial cells RPTEC/TERT1 (HPTEC) were obtained from the American Type Culture Collection (ATCC®; #CRL-4031). These were cultured in DMEM/F12, GlutaMAX supplement (Dulbecco's modified Eagle's medium 1:1 (v/v))

(Life technologies, MA, USA) supplemented with 20 mM Hepes, 2% (v/v) fetal bovine serum (FBS) (HyClone Laboratories, Logan, UT), 5 $\mu$g/ml Apo-transferrin (Sigma-Aldrich, St. Louis, MO, USA), 5 $\mu$g/ml Human Insulin Solution (Sigma), 50 nM dexamethasone (Sigma), 3 nM 3,3',5-Triiodo-L-thyronine sodium salt (Sigma), 10 ng/ml EGF (Sigma), 60 nM Selenium (Sigma), 50 units/ml penicillin and 50 $\mu$g/ml streptomycin (Gibco, Rockville, MD, USA) at 37ºC and 5% $CO_2$. Sub-culturing was performed every 3-5 days. Human primary renal proximal tubule epithelial cells (HprimPTEC) were obtained from the American Type Culture Collection (ATCC®; #PCS-400-010, respectively). These cells were cultured in Renal epithelial cell basal medium (AT ATCC®; #PCS-400-030) supplemented with the Renal epithelial cell growth kit (ATCC®; #PCS-400-040) that contains the following components: Triiodothyronine (10 nM), rhEGF (10 ng/mL), Hydrocortisone Hemisuccinate (100 ng/mL), rh Insulin (5 $\mu$g/mL), Epinephrine (1 $\mu$M), L-Alanyl-L-Glutamine (2.4 mM), Transferrin (5 $\mu$g/mL), 0.5% (v/v) Fetal bovine serum (FBS) at 37ºC and 5% $CO_2$.

Mouse primary kidney cells (MPKC) were isolated from C57BL6/J wild type mice as follows: mice were sacrificed with $CO_2$ overdose and kidneys were collected after perfusion with cold PBS. The capsule was removed and kidneys were minced and digested in PBS containing 2 mg/mL of Collagenase from Clostridium Histolyticum (Sigma-Aldrich) for 20 minutes at 37ºC with gentle stirring, supernatants were sieved through a 70-µm nylon mesh. The cells were incubated with sterile red blood cell lysis buffer (BioLegend, San Diego, CA, USA) and seeded in 10 cm culture dishes. Cells were cultured in RPMI 1640 (Corning, New York, NY) supplemented with 10% FBS, 20 ng/mL EGF (Sigma, St Louis, MO), 50 units/mL penicillin and 50 µg/mL streptomycin (Gibco, Rockville, MD) at 37ºC and 5% $CO_2$**.** I*n vitro* experiments were performed as described below.

**In vitro experiments**

HPTEC, HPrimPTEC and MPKC were plated on 6- well plates and were left on 2%, 0.5% or 10% FBS fresh media, respectively, to reach 80% confluency. Then, cells were synchronized with 100 nM dexamethasone for 2 hours before being washed twice with 1x PBS and placed in 0.5% FBS fresh media for 24 hours. Cells were treated with human recombinant 10 ng/ml TGF-ß1 (R&D

Systems, Minneapolis, MN, USA) and/or 2.5 μM SB505124 (Sigma- Aldrich, St Louis, MO, USA) for 0 to 24 hours in 0.5% FBS media before analyses. HPTEC cells were transfected using lipofectamine 2000 (Invitrogen, Carlsbad, CA, USA) with the aforementioned plasmid constructions. 3xCAGA-luc vector was used as a positive control, Smad3-pCMV5 vector was used for Smad3 overexpression, pCMV5 vector was used as control (these vectors were kindly provided by Dr. Fernando Rodriguez-Pascual). pRL-CMV vector (Promega Corporation, Madison, WI, USA) containing wild-type renilla luciferase was used for normalization in reporter assays at a 1:30 ratio. Cells were incubated with lipofectamine and the corresponding plasmid for 6h at 37ºC. Subsequently, fresh media containing 0.5% FBS was added to the culture and cells were maintained at 37ºC for 24h until reporter analysis or were then treated with human recombinant 10 ng/ml TGF-ß1 and/or 2.5 μM SB505124 for an additional 24 hours for subsequent mRNA and protein expression analysis.

**Immunofluorescence**

HPTEC cells were fixed in 4% neutral buffered formalin for 10 minutes and permeabilized with 0.1% Triton X-100 in PBS for 10 minutes at room temperature (RT). Next, they were blocked with 1% BSA in PBS for 30 minutes at RT and incubated overnight with the Bmal1 and E-cadherin primary antibodies listed in **Table S4** . Then, the samples were incubated for 1 hour for staining with the fluorochrome-conjugated secondary antibody Anti-mouse Alexa488 (Thermo Scientific). Actin cytoskeletons and nuclei were stained with Phalloidin TRITC and DAPI for 40 and 5 minutes at RT, respectively. The coverslips were mounted on slides using MOWIOL (Calbiochem). Cell fluorescence was visualized by a CoolSnap Fx Monochrome confocal microscope with a 40x/1.3 oil Plan-Neofluar M27 objective (Zeiss).

**Reporter assays**

HPTEC cells were lysed with Passive lysis buffer (PLB) (Promega, Madison, WI, USA) 24 hours after transfection and firefly and renilla luciferase activities were determined using a Dual Luciferase Reporter System (Promega Corporation, Madison, WI, USA) and measured using the Glomax-Multi Detection system (Promega, Madison, WI, USA). In order to correct for transfection efficiency, the luciferase activity was normalized to the renilla luciferase activity. Each experimental condition was assayed in triplicate.

**Western Blotting**. Sections of kidney samples were homogenized and HPTEC cells were washed twice with cold PBS and lysed in RIPA buffer including 20 mM Tris-HCl pH 7.5, 150 mM NaCl, 1% sodium deoxycholate, 1% NP-40, 0.1% SDS, protease inhibitors (Complete, Roche Diagnostics, Mannheim, Germany) and phosphatase inhibitors (Sigma). After normalizing for equal protein concentration, lysates were resuspended in SDS sample buffer before separation by SDS-PAGE and transferred onto nitrocellulose membranes. The membranes were incubated with the antibodies listed in **Table S4**. Images were acquired with the Odyssey Infrared Imaging System (LI-COR Biosciences) and densitometry was performed using the ImageJ 1.52p software (NIH, NY, USA).

**RNA isolation and qPCR.** The RNA from kidneys and the different cell cultures was isolated using a phenol-chloroform RNA extraction protocol. RNA samples were treated with RNase-Free DNase set (Qiagen, Cat 79254) following manufacturer's instructions. iScript™ cDNA Synthesis kit (Bio-Rad, Hercules, CA) was used for the reverse transcription, following manufacturer's instructions. qRT-PCR was performed in triplicates using the iQ™SYBR Green Supermix (Bio-Rad, Hercules, CA*)* and the primers described in **Table S5**. Relative mRNA expression was calculated using the ΔΔCt method[61] and the mRNA levels were normalized to 18S.

**TaqMan gene expression assay.** Expression profile of a set of mouse genes related to fibrosis and metabolism was analyzed by using specific and unique TaqMan probes (**Table S6**). After the extraction of RNA, reverse transcription was carried out with the High-Capacity cDNA Reverse Transcription Kit (Thermo Fisher Scientific). RT-PCR was performed with the TaqMan Master (Thermo Fisher Scientific) in a Toche LightCycler 480 Real-Timer PCR system (AB7900HT). Relative mRNA expression was calculated using the ΔΔCt method and the mRNA levels were normalized to ubiquitin.

**Flow cytometry.** Multiparametric flow cytometry was carried out for the identification of monocyte, macrophage and neutrophil cell populations. To accomplish that, kidneys were collected from mice right after perfusion with 1X PBS. They were minced with a blade and incubated in agitation at 37ºC for 20 minutes with 1 mg/ml of collagenase from Clostridium Histolyticum (Sigma-Aldrich) in 1X PBS for tissue disaggregation. Samples were filtered (40 μm) and red blood cells were lysed by their incubation with RBC lysis buffer (BioLegend, San Diego, CA, USA) for 1 minute. Samples were washed twice, centrifuged for 10 min, 2500 rpm at 4ºC and cells were resuspended in 1x PBS. $2x10^6$ cells were pre-incubated with CD16/CD32 (clone 2.4G2, BD, Bioscience, San José, CA, USA) for 5 min at 4ºC in 1X PBS. After two washing cycles with 1x PBS, cells were incubated with Ghost Dye™ Red 780 (Tonbo biosciences, San Diego, CA, USA) for Clock$^{\Delta19/\Delta19}$ mice experiments and Zombie violet™ (BD Biosciences) for CDKO experiment in protein/serum-free 1X PBS for 30 min at 4ºC in darkness to stain dead cells. Cells were washed twice in fluorescence-activated cell sorting (FACS) buffer containing 1% FBS and 0.5% BSA in 1x PBS and then incubated with the antibodies (BioLegend) listed in **Table S7** in darkness for 1h at 4ºC. Antibody incubation was performed in Brilliant Stain buffer (BD Biosciences) for a better resolution of the brilliant violet-conjugated antibodies used in this study. Finally, cells were washed twice and fixed with 4% PFA in 1X PBS. Samples were analyzed on a FACSCanto™ II cytometer with DIVA software (BD Biosciences). Single Stain (SS) and flow minus one (FMO) controls for each fluorophore were used to compensate and establish gates, respectively. Data analysis was performed using FlowJo™ v10.6.2 software (FlowJo, LLC, Ashland, OR, USA). Total cells were gated based in the Forward versus Side scatter (FSC vs SSC). Subsequently, single cells were selected by using Forward – height versus Forward – area (FSC-H vs FSC-A) plot and dead cells were excluded with the corresponding viability marker described above. The identification of inflammatory and hematopoietic cells was based on the presence of CD45. Macrophages were identified by their expression of both F4/80 and CD11b markers. CD86 and CD206 markers were used to identify pro-inflammatory and anti-inflammatory macrophages, respectively. Pro-inflammatory monocytes and neutrophils were identified by their expression of CD11b and the absence of F4/80, and both populations were differentiated by their differential expression of Ly6C and Ly6G.

**Mitochondrial copy number determination.** DNeasy Blood & Tissue Kit (Qiagen Cat. 69504) was used for the extraction of genomic DNA from kidneys according to the manufacturer's instructions. Mitochondrial DNA copy number was characterized with the Mouse Mitochondrial DNA Copy Number Assay Kit (Detroit R&D). Relative mtDNA copy number was represented as the mtDNA to nuclear DNA ratio.

**Soluble collagen quantification.** 5 mg of frozen kidneys were incubated overnight in 0.5M acetic acid with 0.1 mg/mL pepsin (Sigma-Aldrich) at 4ºC. Collagen quantification was performed with Soluble Collagen Assay Kit (Biovision, Cat K532-100) following manufacturer's instructions. Fluorescence was recorded using the ClarioStar plus (Bmg Labtech) and data were normalized with total protein measured with Pierce BCA protein assay kit (Thermo scientific, Cat 23227).

Table S1: Primer sequences for the genotyping of clock mutant mice.

| Name | Sequence (5'-3') | Source |
|---|---|---|
| Clock_wt_Fwd | GGTCAAGGGCTACAGGTA | Erik D. Herzog et al, 1998 |
| Clock_wt_Rv | TGGGGTAAAAAGACCTCTTGCC | Erik D. Herzog et al, 1998 |
| ClockΔ19/Δ19_Fwd | AGCACCTTCCTTTGCAGTTCG | Erik D. Herzog et al, 1998 |
| ClockΔ19/Δ19_Rv | TGTGCTCAGACAGAATAAGTA | Erik D. Herzog et al, 1998 |
| Per2_Fwd | CTGTGTTTACTGCGAGAGT | Seung-Hee Yoo et al, 2004 |
| Per2_wt_Rv | GGGTCCATGTGATTAGAAAC | Seung-Hee Yoo et al, 2004 |
| Per2::Luc_Rv | TAAAACCGGGAGGTAGATGAGA | Seung-Hee Yoo et al, 2004 |
| Cry1_WT_Fwd | GATTCACTGTGTGTTGAGCAAGG | Vitaterna et al, 1999; Vollmers et al., 2012 |
| Cry1_Rev | CTTGAGTGAGTGAGTCTGCTG | Vitaterna et al, 1999; Vollmers et al., 2012 |
| Cry2_WT_Fwd | GCAAGACAGGCTTCCCTTGG | Vitaterna et al, 1999; Vollmers et al., 2012 |
| Cry2_Rev | CACACTGAAATCGGCATCCAG | Vitaterna et al, 1999; Vollmers et al., 2012 |
| Cry1/2_Neo_Fwd | CCTTCTATCGCCTTCTTGACGAG | Vitaterna et al, 1999; Vollmers et al., 2012 |
| Bmal1_Fwd | ACTGGAAGTAACTTTATCAAACTG | Ryota Nakazato et al, 2017 |
| Bmal1LoxP/LoxP_Rv | CTGACCAACTTGCTAACAATTA | Ryota Nakazato et al, 2017 |
| Bmal1_WT_Rv | AATCCGCCTGCCTACTGCCTCC | Ryota Nakazato et al, 2017 |
| Pax8-rtTA_Fwd | CCATGTCTAGACTGGACAAGA | Traykova-BrauchM et al, 2008 |
| Pax8-rtTA_Rv | CTCCAGGCCACATATGATTAG | Traykova-BrauchM et al, 2008 |
| LC-1_Fwd | TCGCTGCATTACCGGTCGATGC | Traykova-BrauchM et al, 2008 |
| LC-1_Rv | CCATGAGTGAACGAACCTGGTCG | Traykova-BrauchM et al, 2008 |

Table S2: Summary of the different procedures performed in mice.

| Time point (Days) | Procedure | Molecular clock mouse model |
|---|---|---|
| 3 | UUO | non-mutated C57BL6/J |
| 7 | | |
| 15 | | |
| 25 | | |
| 7 | FAN | |
| 15 | | |
| 25 | ADN | |
| **Time point (Days)** | **Procedure** | **Molecular clock mouse model** |
| 3 | UUO | WT |
| | | $Clock^{\Delta 19}$ |
| 7 | | WT |
| | | $Clock^{\Delta 19}$ |
| 25 | ADN | WT - Veh |
| | | $Clock^{\Delta 19}$ - Veh |
| | | WT - ADN |
| | | $Clock^{\Delta 19}$ - ADN |
| 3 | UUO | WT |
| | | Bmal1 KO |
| 7 | | WT |
| | | Bmal1 KO |
| 3 | UUO | WT |
| | | CDKO |
| | | Cry1 KO |
| | | Cry2 KO |
| | | CD Het |
| 7 | UUO | WT |
| | | Bmal1 cKO |

Table S2 continued

| Number of mice |
|---|
| 6 |
| 6 |
| 6 |
| 6 |
| 6 |
| 6 |
| 6 |
| 6 |
| 5 |
| 5 |
| **Number of mice** |
| 6 |
| 6 |
| 6 |
| 6 |
| 5 |
| 5 |
| 5 |
| 4 |
| 5 |
| 6 |
| 5 |
| 6 |
| 5 |
| 6 |
| 3 |
| 3 |
| 3 |
| 3 |
| 4 |

Table S3. Primer design for the cloning of human Arntl regulatory regions sequences

| Bases | Nº of bases | annealing Tº |
|---|---|---|
| %GC | 60 | **56ºC** |
| Tm (ºC) | 63 | |
| %GC | 55 | |
| Tm (ºC) | 61 | |
| %GC | 42 | **57ºC** |
| Tm (ºC) | 62 | |
| %GC | 48 | |
| Tm (ºC) | 66 | |

### Table S4. Primer sequence for qPCR analysis

| Region | Primer | Restriction enzyme | Sequence |
|---|---|---|---|
| SmEnh | Forward | Sac1 | 5'CATG**GAGCTC**CTTGCTGGGAGCCATAGGCA3' |
| | Reverse | XhoI | 5'GAT**CTCGAG**TTGGCACCTTGGAAGGTTGG3' |
| hARNTL | Forward | BglII | 5'CG**AGATCT**TATTTATGGTGCACTTGTTGGGT3' |
| | Reverse | NcoI | 5'ATTA**CCATGG**TCCGTCCCTGACCTACTTTCT3' |

Table S5: Primary antibodies for western blot and immunofluorescence. O/N: overnight, RT: room temperature

| Protein | Reference | Species | Dilution | Experimental condition |
|---|---|---|---|---|
| **BMAL1** | sc-365645, Santa Cruz Biotechnology | Mouse | 1/1000 | 4ºC O/N |
| **CLOCK** | ab3517, Abcam | Rabbit | 1/1000 | 4ºC O/N |
| **SMA** | sc-32251, Santa Cruz Biotechnology | Mouse | 1/1000 | 4ºC O/N |
| **Fibronectin 1** | F7387, Sigma | Rabbit | 1/1000 | 4ºC O/N |
| **GAPDH** | MAB374, Millipore | Mouse | 1/15000 | 1h RT |
| **ß -actin** | A1978, Sigma-Aldrich | Mouse | 1/15000 | 1h RT |
| **CPT1A** | Ab128568, Abcam | Mouse | 1/1000 | 4ºC O/N |
| **TFAM** | PA5-68789, Thermo Fisher | Rabbit | 1/1000 | 4ºC O/N |
| **COL1A1** | E8F4L, Cell Signaling | Rabbit | 1/1000 | 4ºC O/N |
| **E-Cadherin** | 610182, Bd transduction lab | Rabbit | 1/1000 | 4ºC O/N |

### Table S6. Primer sequence for qPCR analysis

| Gene | Primer Sequence | |
|---|---|---|
| Mouse | Forward | Reverse |
| 18s | CGGCTACCACATCCAAGGAA | GCTGGAATTACCGCGGCT |
| Adgre1 | AGCTCCCATTCCCAGACTTC | TGCCATCAACTCATGATACCCT |
| Mrc1 | ATGGATTGCCCTGAACAGCA | TGTACCGCACCCTCCATCTA |
| Cd68 | GGGGCTCTTGGGAACTACAC | GTACCGTCACAACCTCCCTG |
| Cd86 | CTTACGGAAGCACCCACGAT | CGGCAGATATGCAGTCCCAT |
| Cd80 | TTCACCTGGGAAAAACCCCC | CCCGAAGGTAAGGCTGTTGT |
| Ly6c | ACTGTGCCTGCAACCTTGTC | TTGGCACTCCATAGCACTCG |
| Ly6g | GTCCACTTCTGAAAGACCTTGT | AGGTGGGACCCCAATACAAT |
| Nos2 | AGGGACTGAGCTGTTAGAGAC | GCACTTCTGCTCCAAATCCA |
| Tnfa | TAGCCCACGTCGTAGCAAAC | GCAGCCTTGTCCCTTGAAGA |
| Il1b | TGCCACCTTTTGACAGTGATG | TGATGTGCTGCTGCGAGATT |
| Il6 | AGCCAGAGTCCTTCAGAGAGAT | GAGAGCATTGGAAATTGGGGT |
| Arg1 | ATGGGCAACCTGTGTCCTTT | TCTACGTCTCGCAAGCCAAT |
| Csf1 | GCCTGTGTCCGAACTTTCCA | AGGGGTGGCTTTAGGGTACA |
| Csf2 | CGTTGAATGAAGAGGTAGAAGTCG | ACTTGTGTTTCACAGTCCGT |
| Ifng | ACAGCAAGGCGAAAAAGGATG | TCTTCCCCACCCCGAATCA |
| Cd163 | TGCTGTCACTAACGCTCCTG | TCATTCATGCTCCAGCCGTT |
| Tgfb1 | CTGCTGACCCCCACTGATAC | GTGAGCGCTGAATCGAAAGC |
| Arntl mPre-mRNA set1 | GGGACAGGCCAAAAGTCTGT | AACAGCCATAGAGCACTCGG |
| Arntl mPre-mRNA set2 | CTATGCAGGTGGCTTGTGGT | ACAGACTTTTGGCCTGTCCC |
| Arntl | AATCGCAAGAGGAAAGGCAGT | TTTGTCCCGACGCCTCTTTT |
| Arntl2 | GCTGTACCGTCCCTGTCAAA | AACCGTCCCATAGCCACAAG |
| Clock | GAGGTCGTCCTTCAGCAGTC | TGTGACATGCCTTGTGGAAT |
| Npas2 | GAGGCTCAATTCAAAGCCAGC | TGTTACCAGGGAGCATGGAG |
| Cry1 | CTGGCGTGGAAGTCATCGT | CTGTCCGCCATTGAGTTCTATG |
| Cry2 | A TGTGTTCCCAAGGCTGTTC | CCTCCTTGGCCATCTTCATA |
| Per1 | ATGCTCGCCATCCACAAGA | GCGGAATCGAATGGGAGAAT |
| Per2 | ATTGGGAGGCACAAAGTCAG | CAGTAGCCGGTGGATTTGTT |
| Nr1f1 | CCCCTACTGTTCCTTCACCA | CCAGGTGGGATTTGGATATG |
| Nr1d1 | CCTCCTTCTATAACGGGAGCCC | CCCACACACCTTACACAGTA |
| **Gene** | **Primer Sequence** | |
| **Human** | **Forward** | **Reverse** |
| 18S | GTTTTCTGGGCCTTCGACCT | TGCCCCTTCTCTCAAATGCT |
| FN1 | GTGGCTGAAGACACAAGGAA | CCTGCCATTGTAGGTGAAT |
| COL1A1 | GAACGCGTGTCATCCCTTGT | GAACGAGGTAGTCTTTCAGCAACA |
| Arntl hPre-mRNA set1 | ACATGCAACGCAATGTCCAG | AGGACAATAGAGCCCAGGGT |
| Arntl hPre-mRNA set2 | GGTGAGACCCTGGGCTCTAT | AACATTCTAGGCAAGGGTGGT |
| ARNTL | GGAAAAATAGGCCGAATGAT | TGAGCCTGGCCTGATAGTAG |
| ARNTL2 | CAGCAGAGTGGAAGATGGTGA | GTTTGTCCAGTTTACGCGCC |
| CLOCK | CAGCCAGTGATGTCTCAAGC | ATGCGTGTCCGTTGTTCC |
| NPAS2 | AACCTCGGCAGCACTTTAAC | GGTTCTGACATGGCTGTGTG |
| CRY1 | CTTGATGCAGATTGGAGCAT | CCATTGGGATCTGTTCTCCT |
| CRY2 | AGGAGAACCACGACGAGA | TCCGCTTCACCTTTTTATAC |
| PER1 | CTGCTACAGGCACGTTCAAG | CTCAGGGACCAAGGCTAGTG |
| PER2 | TTGGACAGCGTCATCAGGTA | TCCGCTTATCACTGGACCTT |
| PER3 | GCAGGTCTATGCCAGTGTGA | ACCACCACCATTCGGTTCT |

### Table S6 (continued)

| | | |
|---|---|---|
| **TIM1** | GATAGAGGCCCATTCCTGCAT | GAAGGGCTGGGGAACTTAGAC |
| **NR1F1** | CTATCCCTCCAAGGCACAAG | AACACAAGACTGACGAGCACA |
| **NR1D1** | ACAGAATCGAACTCTGCACTTCT | GGGGAGGGAGGCAGGTATT |

Table S7: TaqMan probes used for the gene expression profiling.

| TaqMan probes | |
|---|---|
| Gene | Reference |
| **Mitochondria related transcripts** | |
| Ndufv2 | Mm01239727_m1 |
| LrpprC | Mm00511512_m1 |
| Tfam | Mm00447485_m1 |
| Hspa9 | Mm00477716_g1 |
| **Fatty acid oxidation-related genes** | |
| Cpt1a | Mm01231183_m1 |
| Cpt2 | Mm00487205_m1 |
| Acox1 | Mm01246834_m1 |
| Acox2 | Mm00446408_m1 |
| Ppara | Mm00440939_m1 |
| Ppargc1a | Mm01208835_m1 |
| **Fatty acid synthesis:** | |
| Fasn | Mm00662319_m1 |
| Pcx | Mm00500992_m1 |
| Mccc1 | Mm00522414_m1 |
| **Fibrosis-related genes** | |
| col3a1 | Mm00802300_m1 |
| Acta2 | Mm00725412_s1 |
| vim | Mm01333430_m1 |
| Fn | Mm01256744_m1 |
| Col1a1 | Mm00801666_g1 |
| col4a1 | Mm01210125_m1 |
| col2a1 | Mm01309565_m1 |
| Kim-1 | Mm00506686_m1 |
| **Glucose utilization-related genes** | |
| HK | Mm00439344_m1 |
| PFK1 | Mm01309576_m1 |
| PGK1 | Mm00435617_m1 |
| PK | Mm00443090_m1 |
| Pck1 | Mm01247058_m1 |
| G6pc | Mm00839363_m1 |
| Ldh1 | Mm00516030_m1 |
| Ldh2 | Mm00612429_m1 |
| Glut1 | Mm00441480_m1 |
| SDH | Mm00497118_m1 |
| Pkm2 | Mm00834102_gH |
| ATP-synthase and Oxphox | |
| Atp5e | Mm01239887_m1 |
| Ndufs8 | Mm00523063_m1 |
| Cyp4d14 | Mm00484138_m1 |
| **Control genes** | |
| Ubiquitin | Mm01622233_g1 |

Table S8: Fluorochrome-conjugated antibodies for flow cytometry analysis (Biolegend, San Diego, CA, USA)

| Antibody | Reference | Clone | Experiment |
|---|---|---|---|
| **APC anti-human CD45** | 368511 | 2D1 | $Clock^{\Delta19/\Delta19}$ |
| **Percp-Cy5.5 anti-human CD45** | 368505 | 2D1 | CDKO |
| **FITC anti-mouse F4/80** | 123107 | BM8 | $Clock^{\Delta19/\Delta19}$ |
| **APC anti-mouse F4/80** | 123115 | BM8 | CDKO |
| **Percp-Cy5.5 anti-human CD11B** | 301327 | ICRF44 | $Clock^{\Delta19/\Delta19}$ |
| **APC-Cy7 anti-human CD11B** | 301341 | ICRF44 | CDKO |
| **APC anti-mouse CD86** | 105113 | PO3 | $Clock^{\Delta19/\Delta19}$; CDKO |
| **PE anti-mouse CD206** | 141705 | C086C2 | $Clock^{\Delta19/\Delta19}$; CDKO |
| **BV421 anti-mouse Ly6G** | 127627 | 1A8 | $Clock^{\Delta19/\Delta19}$ |
| **BV605 anti-mouse Ly6C** | 128035 | HK1.4 | $Clock^{\Delta19/\Delta19}$ |
| **FITC anti-mouse Ly6G/Ly6C (GR1)** | 108405 | RB6-8C5 | CDKO |

**Supplementary figure legends**

**Figure S1: The gene expression of circadian oscillators is altered in kidney fibrosis.** mRNAexpression of clock-related genes in kidneys from mice subjected to (A) UUO for 3, 7, 15, 25 days (n=6 per condition), data are represented as individual values for obstructed kidneys linked to their corresponding contralateral ones. (B) FAN (n= 6 mice per condition) and (C) ADN (n=5 mice per condition). CTL: Contralateral; Veh: Vehicle. (D) Representative microphotographs of H&E and Sirius red stains from contralateral and obstructed kidneys of WT mice 3, 7, 15 and 25 days after UUO, 15 days after FAN and 25 days after ADN. Scale bar: 100 μm. (E) Quantification of Sirius red in kidney sections of the analysis performed in (D). Data are represented with violin plots as the median with interquartile range. $^{*}$P<0.05, $^{**}$P<0.01 compared to control kidney (black asterisk) or compared to the obstructed kidneys of a different time point for each gene within the same model (grey asterisk). $^{\#}$P<0.05, $^{\#\#}$P<0.01 compared to different time points within the same model, $^{\$\$}$P<0.01 compared to the UUO model.

**Figure S2: The expression of circadian oscillators is upregulated by TGF-β in primary mouse kidney cells and in HprimPTEC.** (A) Relative mRNA levels of Fn1 and clock-related genes in synchronized mouse primary kidney cells treated with TGF-β for 24 hours. Data are represented with violin plots as the median with interquartile range (n=6 per condition) **(B-D)** Evaluation of the expression of circadian oscillators in synchronized human primary proximal tubular epithelial cells treated with TGF-β for 24 hours in the presence or absence of the selective inhibitor SB505124; (B) Relative mRNA levels of Col1a1 and clock-related genes, data are represented with violin plots as the median with interquartile range (n=6 per condition). (C) immunoblots showing the expression of Bmal1, pSmad3, Smad3 and β-actin. β-actin and Smad3 were used for normalization; (D) Relative protein expression of Bmal1and pSmad3 obtained by densitometry of images from C and normalized with β-actin and Smad3, respectively. Data are represented with violin plots as the median with interquartile range (n=6 per condition). $^{\#}$P<0.05, $^{\#\#}$P<0.01 compared to untreated cells, $^{*}$P<0.05, $^{**}$P<0.01 compared to TGF-β-treated cells.

**Figure S3: The expression of the pre-mRNA of Arntl gene is upregulated in kidney fibrosis and by TGF-β in HprimPTEC** (A) Schematic depicting the amplification regions of the primers designed for the analysis of the expression of the mouse (blue) and human (green) pre-mRNA of the gene Arntl. (B) Relative pre-mRNA levels of Arntl in contralateral and obstructed kidneys from mice 25 days after UUO, data are represented with violin plots as the median with interquartile range (n=6 per condition). (C) Relative pre-mRNA levels of Arntl in synchronized human primary proximal tubular epithelial cells treated with TGF-β for 24 hours, data are represented with violin plots as the median with interquartile range (n=4 per condition). $^{\#}$P<0.05, $^{\#\#}$P<0.01 compared to contralateral kidneys or untreated cells.

**Figure S4: *In silico* analysis of the distal enhancer upstream to the human ARNTL gene showing two potential Smad3/4 binding sites and gene expression analysis after Smad3 overexpression.** (A) Representation denoting the regulatory regions located upstream of the ARNTL gene. Data obtained from the UCSC genome browser[56]. The image shows the DNA sequence upstream of the human ARNTL gene and indicates distal and proximal enhancers, H3K27Ac marks and conservation among vertebrates. (B) Matrixes used by JASPAR2018 for the analysis of the potential binding sites. (C) Sites predicted in the distal enhancer by JASPAR2018 with a relative score above 80%. (D)Representation denoting the conservation among vertebrates of the R-Smads potential binding sites found in a distal enhancer close to the ARNTL gene. Data were obtained with the UCSC genome browser[56]. (E) Relative mRNA levels of Col1a1, Arntl and Smad3 in synchronized HPTEC transfected with the overexpression vector for Smad3 (Smad3-PCMV5) or the control vector (PCMV5) and treated with TGF-β for 24 hours in the presence or absence of the selective inhibitor SB505124. Data are represented with box plots as the median with interquartile range (n=6 per condition). (F) Immunoblot showing the expression of Bmal1, pSmad3, Smad3 and β-actin in synchronized HPTEC transfected with the overexpression vector for Smad3 (Smad3-PCMV5) or the control vector (PCMV5) and treated with TGF-β for 24 hours in the presence or absence of the selective inhibitor SB505124. (G) Bioluminescence assays in HPTEC cells transfected with the vector reporter constructions Pgl3p, hARNTLp, SmEnh-Pgl3p, SmEnh-hARNTLp, 3xCAGA and with theoverexpression vector for Smad3 treated with TGF-β for 24 hours in the presence or absence of the selective inhibitor SB505124. Data are represented as the mean ± SEM $^{\#\#}$P<0.01, $^{\#\#\#}$P<0.001 compared to control without treatments, $^{**}$P<0.01 compared to cells treated with TGF-β only. $^{\$}$P<0.05 compared to their corresponding control without Smad3 overexpression.

**Figure S5: Circadian maladjustment in the different circadian oscillator- deficient mouse models.** (A) Mice activity measured in the Bmal1 KO and their corresponding Wt mice for twodays in a regular 12h light:12h dark cycle and 2 days in constant darkness (B-E) mRNA expression of circadian oscillators in normal kidneys of (B) Clock$^{\Delta19}$ mice, (C) Bmal1 KO mice, (D) Bmal1 cKO mice and in contralateral and obstructed kidneys of (E) Cry1/Cry2 knockout mice 3 days afte UUO. Data are represented in a box plot as the median with interquartile range. Number of mice: Clock mutant mice: WT (6) and Clock$^{\Delta19}$ mice (6); Bmal1 KO: WT (5) and Bmal1 KO (6); Bmal1 cKO: WT (3) and Bmal1 cKO (4); Cry-deficient mice: WT (n=5), CDKO (n=6), Cry1 KO (n=3), Cry2 KO(n=3), CDhet mice (n=3). $^{\#}$P<0.05, $^{\#\#}$P<0.01, $^{\#\#\#}$P<0.001 compared to WT; $^{\$}$P<0.05, compared to their respective contralateral kidney.

**Figure S6: Bmal1 deficiency and Clock mutation do not exhibit significant differential impairment in fibrosis 7 days after UUO.** (A) Schematic of experimental design (B) Representative microphotographs of H&E, Sirius red and KIM1 immunohistochemistry (IHC) stains from contralateral and obstructed kidneys of Bmal1 KO, Clock$^{\Delta19}$ and their corresponding WT mice 7 days after UUO. Scale bar: 100 µm. (C, D) Quantification of (C) Sirius red and (D) KIM1 IHC in kidney sections of the different genetically-modified mouse models denoted in (B), data are represented in a box plot as the median with interquartile range. (E) Immunoblot depicting the expression of αSMA in kidneys from the Bmal1 KO mice 7 days after UUO, GAPDH levels were used for normalization. (F) Relative protein expression of αSMA in kidneys from Bmal1 KO mice, data are represented in a box plot as the median with interquartile range. (G) Plasma creatinine and (H) BUN levels of Bmal1 KO mice and their respective WT mice 7 days after UUO. Data are represented in a box plot as the median with interquartile range. (I) Relative mRNA expression of fibrosis-related genes in WT and Bmal1 KO mice 7 days after UUO. Data are represented in a box plot as the median with interquartile range. (J) Immunoblot depicting the expression of αSMA in kidneys from the Clock$^{\Delta 19}$ mice 7 days after UUO, GAPDH levels were used for normalization. (K) Relative protein expression of αSMA in Clock$^{\Delta 19}$ mice of images in J. Data are represented in a box plot as the median with interquartile range. (L) Relative mRNA expression of fibrosis-related genes in WT and Clock$^{\Delta 19}$ mice 7 days after UUO, data are represented in a box plot as the median with interquartile range. (Number of mice: Bmal1 deficient mice: WT (5) and Bmal1 KO (6); Clock$^{\Delta 19}$ mice: WT (6) and Clock mice (6). $^{*}P<0.05$, $^{**}P<0.01$ compared to their corresponding contralateral kidney; $^{\#}P<0.05$, $^{\#\#}P<0.01$ compared to WT.

**Figure S7: Bmal1 cKO mice do not show differentially increased fibrosis 7 days after UUO.** (A) Schematic of experimental design. (B) Representative microphotographs of Sirius red staining from contralateral and obstructed kidneys of WT and Bmal1 cKO mice 7 days after UUO; (C) Quantification of Sirius red staining in kidney sections of mice denoted in (B), the data are represented in a box plot as the median with interquartile range; (D) Relative mRNA expression of fibrosis-related genes in the kidneys from WT and Bmal1 cKO 7 days after UUO; (E) Immunoblot depicting the expression of αSMA. Tubulin levels were used for normalization; (F) Relative protein expression of αSMA obtained by densitometry and normalized with tubulin. Data are represented in a box plot as the median with interquartile range. Number of mice: WT (n=3) and Bmal1 cKO (n=4). $^{*}P<0.05$, $^{***}P<0.01$ compared to their corresponding contralateral kidney.

**Figure S8: Clock$^{\Delta 19}$ mice exhibit an increased presence of anti-inflammatory macrophages during the late phase of inflammation while no significant differences were observed in Bmal1 deficient mice.** (A) Schematic of experimental design of Clock$^{\Delta 19}$ mice subjected to adenine-induced nephropathy. (B) Flow cytometry dot plots denoting the expression of inflammatory-related markers selected for the analysis of different innate immune populations. CD45 is a panleukocyte antigen also present in hematopoietic cells, monocytes and macrophages. F4/80 is a surface marker expressed in monocytes and macrophages and CD11b is a marker commonly expressed in inflammatory myeloid cells. CD86 and CD206 were used for the selection of proinflammatory or anti-inflammatory macrophage subpopulations (CD45+F4/80+CD11b+), respectively. Ly6c and Ly6g were used for the selection of monocytes (CD45+F4/80-CD11b+Ly6c+Ly6g-) and neutrophils (CD45+F4/80-CD11b+Ly6c+Ly6g+) in kidneys from Wt and Clock$^{\Delta 19}$ mice, 25 days after the start of adenineadministration. (C, D) Quantification of the different immune populations analyzed in (B), data are represented in a box plot as the median with interquartile range. (E) Schematic of experimental design of Clock$^{\Delta 19}$ mice subjected to UUO for 7 days. (F, G) Relative mRNA expression of inflammatory-related genes in kidneys from Clock$^{\Delta 19}$ mice 7 days after UUO. Data are represented in a box plot as the median with interquartile range. (H) Schematic of experimental design of Bmal1 KO mice subjected to UUO for 7 days. (I, J) Relative mRNA expression of inflammatory-related genes in kidneys from Bmal1 KO mice 7 days after UUO, data are represented in a box plot as the median with interquartile range. (K) Schematic of experimental design of Bmal1 cKO mice subjected to UUO for 7 days. (L) Relative mRNA expression of inflammatory-related genes in kidneys from Bmal1 cKO mice 7 days after UUO, data are represented in a box plot as the median with interquartile range. Number of mice: Clock$^{\Delta 19}$: WT Veh (5), WT ADN (5), WT UUO (6), Clock$^{\Delta 19}$ Veh (5), Clock$^{\Delta 19}$ ADN (4), Clock$^{\Delta 19}$ UUO (6); Bmal1 deficient mice: WT (5), Bmal1 KO (6); Bmal1 conditional KO: WT (3), Bmal1 cKO (4). $^{*}P<0.05$, $^{**}P<0.01$ $^{***}P<0.001$ compared to their respective contralateral kidney; $^{\#}P<0.05$ compared to WT.

**Figure S9: The expression profile of metabolism and mitochondrial-related genes is significantly altered in Bmal1 KO mice while no significant differences were observed in Bmal1 cKO and Clock$^{\Delta 19}$ mice.** (A) Schematic of experimental design of Bmal1 KO mice subjected to UUO for 3 days. (B) Heat map of normalized expressions of metabolism and mitochondrial-related genes in kidneys from WT and Bmal1 KO mice, 3 days after UUO. Yellow rectangles denote significant differences compared with control kidneys from Wt mice. (C) Immunoblot depicting the expression of Cpt1a in the kidneys from Bmal1 KO mice 3 days after UUO; GADPH levels were used for normalization. (D) Relative protein expression of Cpt1a in the kidneys from Bmal1 KO mice of images in C; data are represented in a box plot as the median with interquartile range. (E) Schematic of experimental design of Clock$^{\Delta 19}$ mice subjected to UUO for 3 days. (F) Heat map of normalized metabolism and mitochondrial-related genes in Clock$^{\Delta 19}$ mice

7 days after UUO. (G) Immunoblot depicting the expression of Cpt1a and TFAM in the kidneys of Clock$^{\Delta 19}$ mice; GADPH levels were used for normalization. (H) Relative protein expression of Cpt1a in the kidneys of Clock$^{\Delta 19}$ mice of images in G, data are represented in a box plot as the median with interquartile range. (I) Schematic of experimental design of Bmal1 cKO mice subjected to UUO for 7 days. (J) Heat map of normalized metabolism and mitochondrial-related genes in Bmal1 cKO mice 7 days after UUO. (K) Immunoblot depicting the expression of Cpt1a a TFAM in the kidneys of Bmal1 cKO mice; GADPH levels were used for normalization. (L) Relative protein expression of Cpt1a in the kidneys of Bmal1 cKO mice of images in K, data are represented in a box plot as the median with interquartile range. Number of mice: Bmal1 deficient mice: WT (5) and Bmal1 KO (6); Clock deficient mice: WT (6) and Clock$^{\Delta 19}$ mice (6); Bmal1 conditional KO: WT (3) and Bmal1 cKO (4). $^{**}P<0.01$ compared to their respective contralateral kidney.
